# Capsid lattice destabilization leads to premature loss of the viral genome and integrase enzyme during HIV-1 infection

**DOI:** 10.1101/2020.03.26.009902

**Authors:** Jenna E. Eschbach, Jennifer L. Elliott, Wen Li, Kaneil K. Zadrozny, Keanu Davis, Shawn Mohammed, Dana Q. Lawson, Owen Pornillos, Alan N. Engelman, Sebla B. Kutluay

## Abstract

The human immunodeficiency virus type 1 (HIV-1) capsid (CA) protein forms a conical lattice around the viral ribonucleoprotein complex (vRNP) consisting of a dimeric viral genome and associated proteins, together constituting the viral core. Upon entry into target cells, the viral core undergoes a process termed uncoating, during which CA molecules are shed from the lattice. Although the timing and degree of uncoating are important for reverse transcription and integration, the molecular basis of this phenomenon remains unclear. Using complementary approaches, we assessed the impact of core destabilization on the intrinsic stability of the CA lattice in vitro and fates of viral core components in infected cells. We found that substitutions in CA can impact the intrinsic stability of the CA lattice in vitro in the absence of vRNPs, which mirrored findings from assessment of CA stability in virions. Altering CA stability tended to increase the propensity to form morphologically aberrant particles, in which the vRNPs were mislocalized between the CA lattice and the viral lipid envelope. Importantly, destabilization of the CA lattice led to premature dissociation of CA from vRNPs in target cells, which was accompanied by proteasomal-independent losses of the viral genome and integrase enzyme. Overall, our studies show that the CA lattice protects the vRNP from untimely degradation in target cells and provide the mechanistic basis of how CA stability influences reverse transcription.

**AUTHOR SUMMARY:** The human immunodeficiency virus type 1 (HIV-1) capsid (CA) protein forms a conical lattice around the viral RNA genome and the associated viral enzymes and proteins, together constituting the viral core. Upon infection of a new cell, viral cores are released into the cytoplasm where they undergo a process termed “uncoating”, i.e. shedding of CA molecules from the conical lattice. Although proper and timely uncoating has been shown to be important for reverse transcription, the molecular mechanisms that link these two events remain poorly understood. In this study, we show that destabilization of the CA lattice leads to premature dissociation of CA from viral cores, which exposes the viral genome and the integrase enzyme for degradation in target cells. Thus, our studies demonstrate that the CA lattice protects the viral ribonucleoprotein complexes from untimely degradation in target cells and provide the first causal link between how CA stability affects reverse transcription.

## INTRODUCTION

Formation of infectious HIV-1 virions is coordinated by the major structural polyproteins Gag and Gag-Pol. Gag selectively packages a dimeric viral genome, targets particle assembly to the plasma membrane, and oligomerizes with other Gag and Gag-Pol polyproteins at the plasma membrane primarily through interactions between the capsid (CA) domains of neighboring Gag molecules [1, 2]. Following the budding of immature virions, the virally encoded protease enzyme cleaves Gag and Gag-Pol polyproteins into their constituent domains triggering virion maturation [1, 2]. Virions undergo a major structural rearrangement, such that the cleaved CA monomers form a conical lattice in which the viral genome condenses with both the cleaved nucleocapsid (NC) domain of Gag and the Pol-encoded viral enzymes, reverse transcriptase (RT) and integrase (IN), to form the viral core [3].

The mature HIV-1 core contains ~250 hexameric and 12 pentameric rings of CA that are stabilized through an extensive network of intra- and inter-subunit interactions between CA molecules [4–9]. Within pentamers and hexamers, the N-terminal domain (NTD) of one CA molecule interacts with a groove in the C-terminal domain (CTD) of the neighboring CA molecule. The first three helices of the NTD interact to form an 18-helix bundle (or 15-helix bundle for pentamers) at the center of the hexamer. Inter-hexamer connections forming the hexagonal lattice are mediated through CTD-CTD interactions. In addition, recent studies revealed that a small molecule, inositol hexakisphosphate (IP_6_), can facilitate the assembly of the CA lattice [10] and regulate its stability [11]. Mutations or compounds that target the critical interactions between individual CA subunits disrupt processes ranging from particle assembly, virion morphogenesis, reverse transcription, and nuclear entry in target cells, underscoring a wide range of functional requirements for the CA protein and/or capsid lattice in multiple steps of the viral life cycle [12–17].

Following their release into the cytoplasm of target cells, HIV-1 cores undergo a poorly understood process termed uncoating, i.e. shedding of CA subunits from the core. The current consensus in the field is that viral cores undergo various levels or stages of uncoating [18–20]. First, a large amount of virion-associated CA appears to be lost soon after entry [21–25]. This is likely due to a combination of uncoating as a result of the metastable structure of the CA lattice and dispersal of CA molecules that are incorporated into virions but are not part of the CA lattice [26–29]. A second phase of uncoating takes place during or as a result of reverse transcription [21, 23, 25, 30–32]. Additionally, a number of cellular proteins that bind CA have been proposed to regulate core stability and uncoating [33]. Although the majority of virion-associated CA is lost during uncoating, both biochemical and genetic evidence supports the notion that some CA remains associated with the reverse transcription complex (RTC) and pre-integration complex (PIC) that respectively mediate reverse transcription and integration during virus infection: CA is the major determinant for HIV-1 nuclear entry [34–41], a fraction of CA remains physically associated with the PIC [42–47], CA contributes to viral DNA (vDNA) integration into actively transcribed genes [39, 48—51], and CA may influence innate host responses by shielding the reverse transcription products from cGAS-STING-mediated sensing [52–56].

Proper uncoating of the HIV-1 core and reverse transcription appear to be interconnected processes. Mutations in CA that destabilize the core in vitro block reverse transcription in target cells [13, 57–59]. Additionally, reverse transcription can accelerate or, if inhibited, delay the uncoating of the CA lattice [30, 31, 60–62]. Exactly how altering the stability of the CA lattice causes defects in reverse transcription is unclear; however, the underlying mechanism may be similar to that which leads to the reverse transcription defects observed upon inhibition of HIV-1 integrase (IN)-RNA interactions.

The HIV-1 IN enzyme has recently been shown to carry out a non-catalytic role in particle maturation through its binding to the viral RNA (vRNA) genome [63]. Inhibition of IN-RNA interactions yields morphologically aberrant particles in which the vRNPs composed of the vRNA and associated enzymes are mislocalized outside the CA lattice [63–65]. Much like viruses with altered core stability, these viruses are blocked at an early reverse transcription stage in target cells [63–89], which can be explained partly by the premature degradation of the unprotected vRNA [90]. Curiously, viruses generated in the presence of a CA-targeting compound, C1, also yield morphologically aberrant particles that are blocked at reverse transcription [91]. Whether CA destabilization affects IN-RNA interactions and whether degradation of the unprotected vRNPs underlies the reverse transcription defect upon CA destabilization remains unexplored.

HIV-1 uncoating has been a difficult process to study due to the metastable nature of the CA lattice and relatively high particle-to-infectivity ratio of HIV-1 preparations that indicate that the vast majority of virus particles are non-infectious [19]. Biochemical and microscopy-based approaches are the current standard and have been widely utilized in the field. Recently, a reporter assay system exploiting the cytoplasmic exposure of a virion-associated mRNA was reported [92]. Previous work that identified key mutations in CA important for core stability [13, 16, 93] depended solely on an in vitro core disassembly assay [13, 93]. While the increased rate of core disassembly in this system correlated with reverse transcription defects in cells, it is untested whether core disassembly also occurs in the context of cell infection. Microscopy-based experiments partially fill this gap and can provide single-cell level information about the kinetics of the early stages of virus replication [21, 22, 24, 25]. Such approaches are generally limited by the difficulty in distinguishing infectious from non-infectious virus particles, albeit more elaborate live-cell imaging approaches have recently been developed to address this shortcoming [22, 25]. Another limitation of microscopy-based approaches is their dependence on indirect labeling of core components. Biochemical separation of post-nuclear supernatants from infected cells, referred to as fate of capsid/core assay, addresses some of these shortcomings and provides an easily accessible alternative [90, 94, 95]. The main advantage of this approach is the ability to trace virtually every component of the HIV-1 core [90, 95], and bypass potential artifacts due to indirect labeling of CA or use of fusion proteins. However, as this approach is laborious and has inherent limitations due to the analysis of bulk cell lysates, it has not been widely adopted to study the effects of CA stability on core components in infected cells. Given the discrepancies between microscopy-based and biochemical approaches [21, 23, 30], and the pros and cons of each approach, it is advantageous to utilize complementary assays to study early post-entry events in the HIV-1 life cycle.

Here, we took an in depth approach to examine the effects of widely utilized CA stabilizing/destabilizing mutations and a CA targeting compound, C1 [91, 96, 97], on the physical properties of the CA lattice, virion architecture, and fates of core components in target cells. We found that CA destabilizing substitutions (P38A, K203A, Q219A) significantly decreased and a CA stabilizing E45A substitution increased the intrinsic stability of the CA lattice. Unstable CA mutants tended to increase the propensity to form eccentric particles with vRNPs mislocalized between the empty CA lattice and the lipid envelope without impacting IN-RNA interactions. Most notably, we found that CA destabilizing mutations and C1 led to dissociation of CA from vRNPs in target cells, which was accompanied by the premature loss of the vRNA and the IN enzyme. Overall, our studies show that the CA lattice protects the viral core components from untimely degradation in target cells and provide the long sought causal link between core stability and reverse transcription.

## RESULTS

### Effects of CA stability on general properties of HIV-1 virions

We first assessed the effects of CA substitutions that were previously reported to decrease (i.e. P38A, Q63A/Q67A, K203A, Q219A) or increase (i.e. E45A) the stability of the CA lattice [13, 16] on HIV-1 replication. The locations of the targeted amino acid residues are dispersed throughout the hexameric CA structure, positioned at the NTD-NTD interface (Pro38 and Glu45) or NTD-CTD interface (Gln63A and Gln67) within the hexamer [4], or at the 3-fold CTD-CTD interhexameric interface (Lys203A and Gln219) [12] (Fig. 1A). Missense mutations were introduced into the replication-competent pNL4-3 molecular clone or NL4-3-derived Gag-Pol expression plasmid for use in subsequent assays described below. With the exception of the Q63A/Q67A substitutions, which substantially impaired particle release, none of the other changes measurably affected Gag expression, processing, or particle release (Fig. 1B, D). All substitutions decreased virus titers circa 10− to >100-fold (Fig. 1C, E), as expected from previous observations [13, 16]. In parallel, we assessed the effects of Compound 1 (C1) [91] on HIV-1 replication. Addition of C1 to virus-producing cells decreased virus titers 10-20 fold (Fig. 1F) without impacting Gag expression, processing (Fig. 1G, H), or particle release (Fig. 1H). As previously noted [91], we observed a dose responsive decrease in the levels of unprocessed Gag in virions, without any corresponding change in processed CA levels in virions (Fig. 1H).

**Figure 1.**
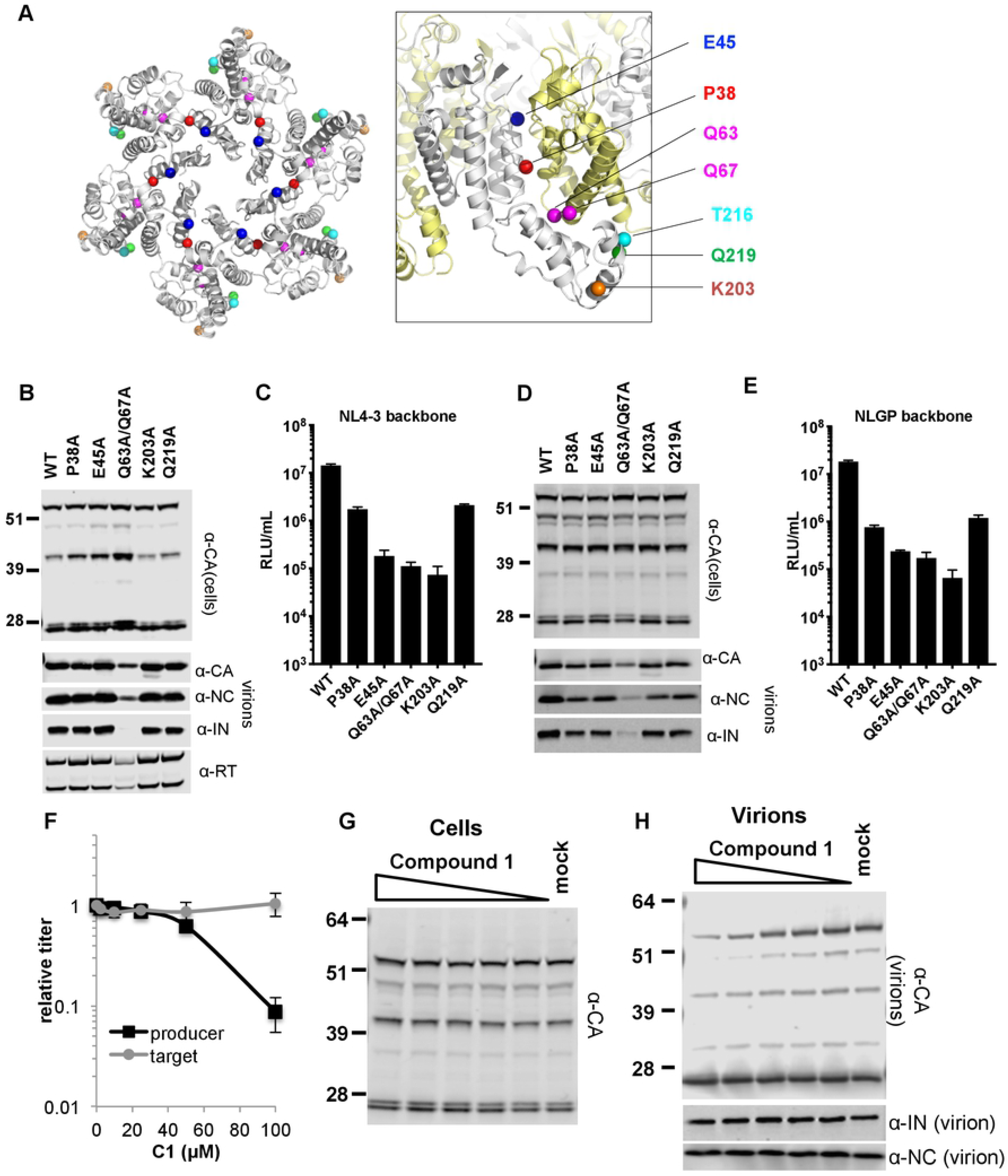
CA stability mutants and compound 1 (C1)-treated viruses. **(A)** Location of CA mutations depicted on the top view of the CA hexamer crystal structure (PDB 4XFX [144]). Close-up of two adjacent CA subunits, colored in gray and yellow with substituted residues shown as colored spheres, is depicted on the right. **(B, C)** HEK293T cells were transfected with pNL4-3 bearing the indicated CA mutations. Cell lysates and purified virions were harvested two days post transfection and analyzed by immunoblotting for CA, NC, IN and RT (B). Virus titers were determined on TZM-bl cells (C). **(D, E)** HEK293T cells were transfected with NLGP-expression plasmid bearing the indicated CA mutations alongside a packagable vRNA (V1B) and VSV-G expression plasmid. Cell lysates and purified virions were harvested two days post transfection and analyzed by immunoblotting for CA, NC and IN (D), and virus titers were determined on TZM-bl cells (E). **(F-H)** HEK293T cells transfected with pNL4-3 were treated with indicated concentrations of C1. Resulting virus titers were determined using TZM-bl cells (F). Cell lysates (G) and purified virions (H) were harvested two days post transfection and analyzed by immunoblotting for CA, NC and IN. Data in A-H is representative of 2-4 independent experiments.

### Effects of CA mutations on virion morphology and IN-RNA interactions

Qualitative assessment of CA destabilizing substitutions on virion morphology has previously revealed the presence of mature particles with fully formed CA cones [13, 16]. In contrast, CA destabilization by C1 increased the occurrence of eccentric particles in which vRNPs were mislocalized outside of the CA lattice [91]. Given these seemingly opposite effects, we wanted to reassess quantitatively the impact of CA destabilizing substitutions on virion morphology. As expected, thin-section electron microscopic (TEM) analysis of cell-free wild-type (WT) HIV-1 particles revealed that ~85% of the virions contained conical cores with centrally-located electron dense vRNPs (Fig. 2A, B). The remaining virions were classified as having either an immature or eccentric morphology with vRNPs localized between the viral membrane and an electron-lucent core (Fig. 2A, B). Most of the CA substitutions including P38A, E45A, Q63A/Q67A, and K203A significantly increased the occurrence of eccentric particles (Fig. 2A, B). The Q219A substitution also marginally enhanced the occurrence of the eccentric particle morphology (Fig. 2A, B).

**Figure 2.**
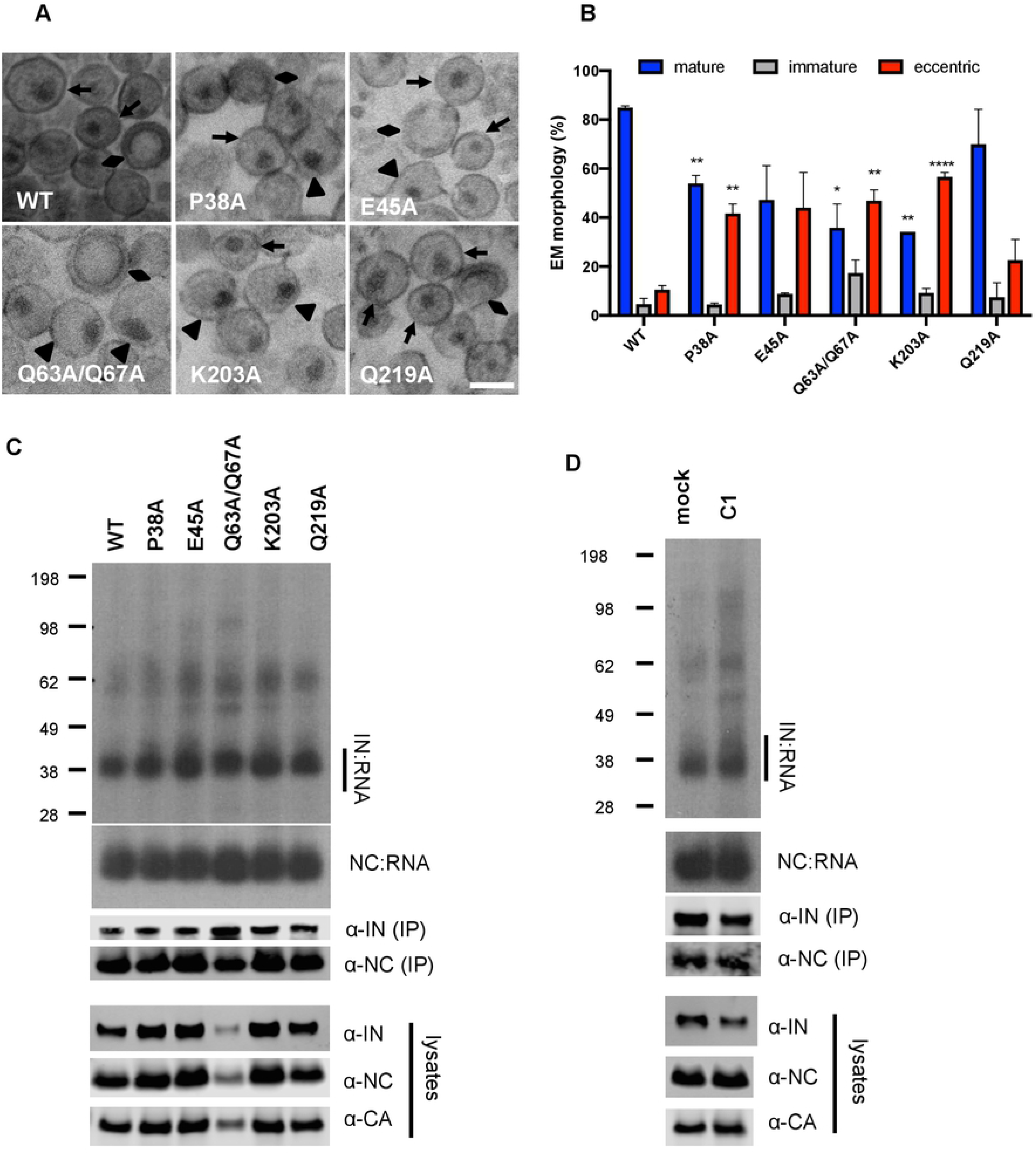
CA stability mutants alter particle morphology but not IN-RNA and NC-RNA complex formation. **(A)** Representative TEM images of WT and CA mutant HIV-1_NL4-3_ virions. Magnification is 30,000x (scale bar, 100 nm). Black arrows indicate mature particles containing conical or round cores with associated electron density; triangles indicate eccentric particles with electron dense material situated between translucent cores and the viral membrane; diamonds indicate immature particles. **(B)** Quantification of virion morphologies from two independent experiments (average ± SD; more than 100 particles counted per experiment). Significance of differences versus matched WT morphology was assessed by t-test. p<0.05 (*), p<0.01 (**), p<0.001 (***), p<0.0001(****). **(C, D)** Representative autoradiograms of IN-RNA and NC-RNA adducts immunoprecipitated from WT or CA mutant HIV-1_NL4-3_ virions (C) or HIV-1_NL4-3_ virions generated from cells treated with 100 μM C1 (D). The amount of immunoprecipitated material was normalized such that equivalent levels of WT and mutant IN proteins were loaded on the gel, as also evident in the immunoblots shown below. Levels of IN, NC and CA in input virion lysates is shown in the lower immunoblots. Data are representative of three independent replicates.

Because loss of IN binding to the viral genome significantly extenuates eccentric particle morphology [63–65], we next investigated the extent of IN-RNA interactions within the different CA mutant viruses. IN-RNA complexes were immunoprecipitated from UV-crosslinked virions and the amount of vRNA bound by IN was assessed by end-labeling followed by separation of protein-RNA complexes by gel electrophoresis [63, 98, 99]. Equivalent levels of IN-RNA complexes were isolated from WT virions and viruses bearing CA substitutions (Fig. 2C) or WT virus generated in the presence of C1 (Fig. 2D). Likewise, CA substitutions and C1 treatment did not seem to impact NC-RNA interactions in virions (Fig. 2C, D).

### Effects of CA substitutions on the intrinsic stability of the CA lattice

The traditional approach to assess CA stability is based on isolation of cores from envelope-stripped virions followed by equilibrium density sedimentation on linear sucrose gradients, during which the cores migrate to denser fractions [13]. The fraction of CA in the dense fractions is assumed to directly correlate with the extent of uncoating, but the assay does not distinguish between contributions to overall core yield that arise from the intrinsic stability of the capsid lattice versus modulatory effects of other factors. To assess the impact of CA mutations on the intrinsic stability of the mature CA lattice, WT CA and CA proteins bearing the aforementioned substitutions were assembled into capsid-like tubes in vitro in the absence of any other viral or cellular factors, diluted, and analyzed by nano-differential scanning fluorimetry (nano-DSF) during thermal denaturation (Fig. 3A). Dilution of assembled tubes in this assay results in their partial disassembly, and we reasoned that the degree of disassembly would reflect the intrinsic stability of the CA structures. Given the low yield of particles obtained with CA mutant Q63A/Q67A (Fig 1B, D & Fig. 2C), this mutant was excluded from the remainder of the studies described below.

**Figure 3.**
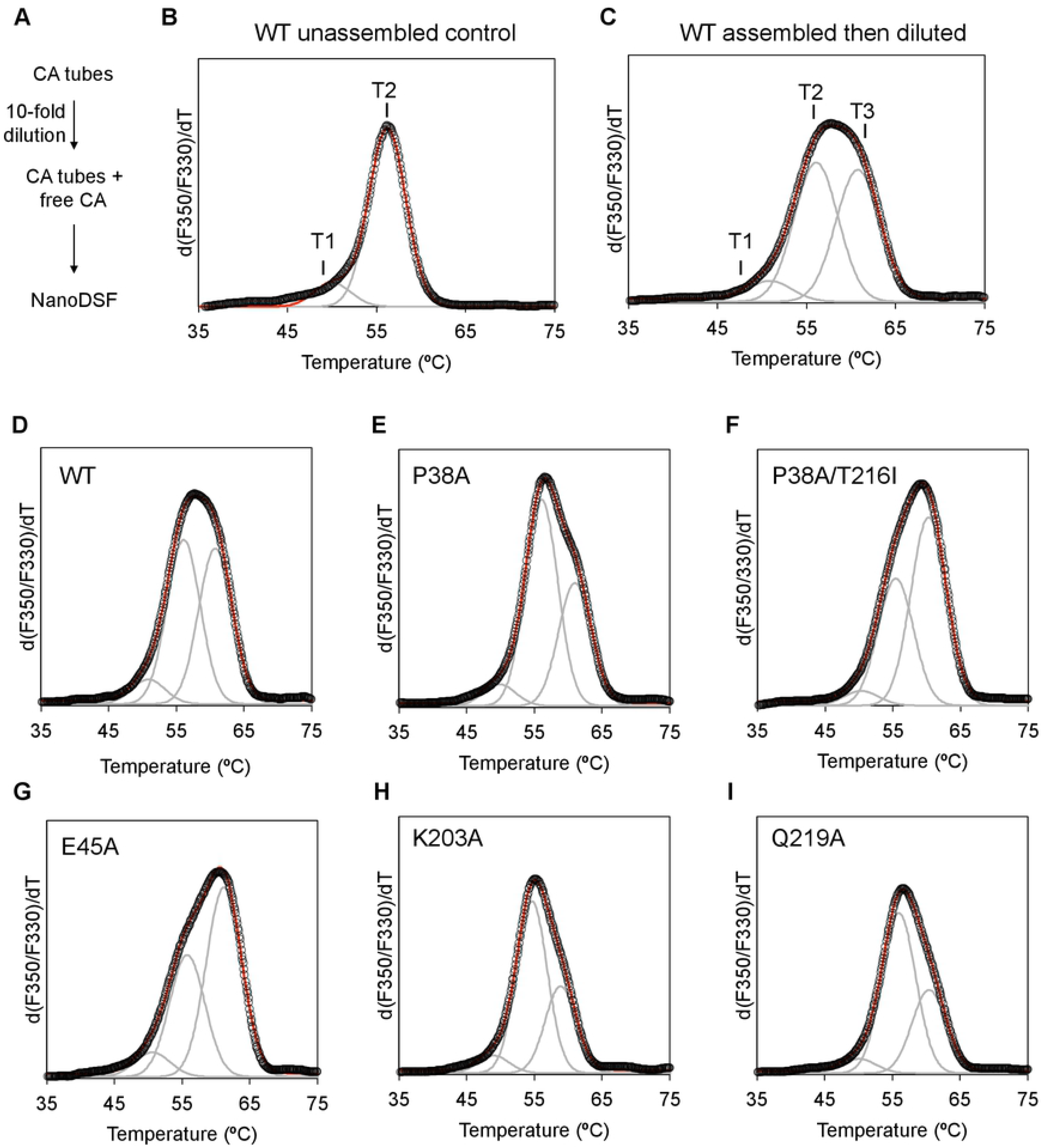
Estimation of relative intrinsic CA tube stability. (A) Schematic of the nanoDSF assay. Assembled tubes were diluted 10-fold into assembly buffer and the thermal melting profile of the sample was measured. (B) The first derivative profile (black circles) of unassembled WT protein control can be deconvoluted into two component peaks of distinct melting points, indicated by T1 and T2. The components, which we assign to the NTD (T2) and CTD (T1), are modeled as Gaussian curves (gray). The sum of the Gaussian components is shown by the red curve. (C) The profile of assembled WT CA tubes contains an additional, third component with a higher melting point (T3), which arises from the fraction of protein that remained assembled after dilution. The height of the T3 Gaussian component compared to T2 and T1 indicates the relative proportion of intact tubes that remained after dilution, and is therefore a direct readout of intrinsic CA lattice stability. (D-I) Profiles of the indicated mutants. Data is representative of two independent biological replicates.

Comparison of unassembled WT CA (Fig. 3B) and assembled CA tubes (Fig. 3C, D) revealed the presence of the T3 species with greater thermal stability than initial T1 and T2 populations following assembly (Fig. 3C, D). The P38A (Fig. 3E), K203A (Fig. 3H), and Q219A (Fig. 3I) substitutions each decreased the levels of the stable T3 species as evident by the skewing of the nano-DSF curves towards the left, consistent with a destabilizing effect on the capsid lattice. The opposite was true for the E45A substitution, which showed an increased proportion of the more stable species (Fig. 3G); this result is indicative of lattice stabilization and is in line with previous observations [100]. Adding the T216I substitution (see Fig. 1 for Thr216 location) to P38A, which partially restored the P38A infectivity defect [101], similarly increased CA tube stability (Fig. 3F). Together, these results provide direct evidence that CA mutations can impact the intrinsic stability of the assembled CA lattice in the absence of other virion and core components.

### Effects of CA mutations on core stability in vitro

We next validated how modulation of CA alters core stability in virions by utilizing a biochemical assay in which the viral lipid envelope is stripped off by brief detergent treatment and core components are separated by equilibrium density sedimentation on linear sucrose cushions [90, 102]. Note that while similar approaches have been previously employed to study the stability of isolated CA mutants [13, 91, 103, 104], a side-by-side comparison of the effects of multiple CA substitutions on the behavior of different core components was lacking.

Following centrifugation of envelope-stripped virions, fractions collected from the top of the gradients were analyzed for the presence of CA, IN, MA, and RT activity. For WT viruses, a large fraction of CA migrated in top fractions representing soluble CA that has dissociated from the CA lattice during the assay and/or CA that was incorporated into virions but was not part of the CA lattice [26–28] (Fig. 4A, B). A second population of CA was present in dense fractions 7-10, representing CA that is in complex with dense vRNPs. The P38A, K203A, and Q219A substitutions as well as treatment with C1 each led to a substantial decrease in the levels of CA in dense fractions, whereas the E45A substitution tended to yield modestly higher levels of CA compared to WT viruses (Fig. 4A, B, S1A). As anticipated, MA remained primarily in soluble fractions confirming the efficient removal of the viral lipid envelope (Fig. 4C). RT activity traces mirrored those of CA protein, with 10-50-fold less RT activity in dense fractions upon destabilization of the CA lattice (Fig. 4D, S1B). IN (Fig. 4E, F) and vRNA (not shown) remained in dense fractions for all viruses, suggesting that CA mutations and C1 did not alter vRNP condensation, consistent the IN-RNA and NC-RNA binding profiles (Fig. 2C, D).

**Figure 4.**
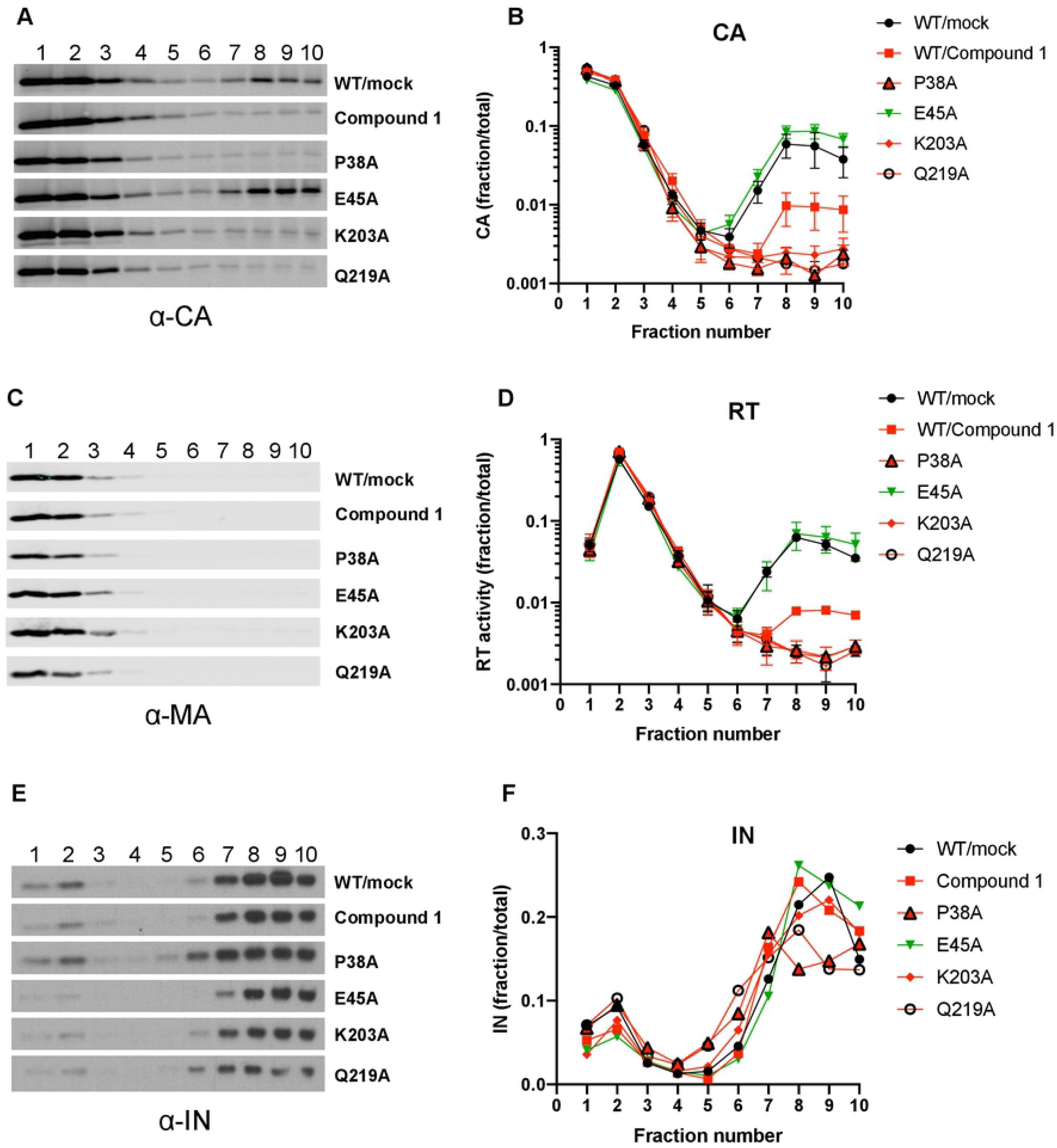
Effects of CA mutations and C1 on the stability of HIV-1 cores in vitro. HIV-1_NL4-3-_derived virions bearing the indicated CA mutations or grown in the presence of 100 μM C1 were subjected to equilibrium density sedimentation following treatment with 0.5% Triton as detailed in Materials & Methods. Ten fractions collected from the top of the sucrose gradients were analyzed for CA (A, B), MA (C), RT (D), and IN (E, F) by immunoblotting or a Q-PCR-based RT activity assay. Results in B & D (mean values ± SEM) quantify CA content and RT activity from three independent biological replicate experiments; panel F quantifies the results of one of three representative IN immunoblots.

In contrast with its stabilizing effect on CA tubes (Fig. 3F), the T216I substitution that partially compensated the P38A infectivity defect did not counteract the loss of CA (Fig. 5A, B, S2A) or RT (Fig. 5C, S2B) in dense fractions, consistent with a previous study [103]. In contrast, the R132T substitution that conferred resistance to C1 significantly increased the amount of CA (Fig. 5D, E, S2C) and RT (Fig. 5F, S2D) in dense fractions. As above, IN remained in dense fractions under all conditions (Fig. 5A, 5D). Overall, these results indicate that decreasing the stability of the CA lattice can lead to dissociation of CA and RT from vRNPs, likely without impacting condensation of vRNPs by NC and IN.

**Figure 5.**
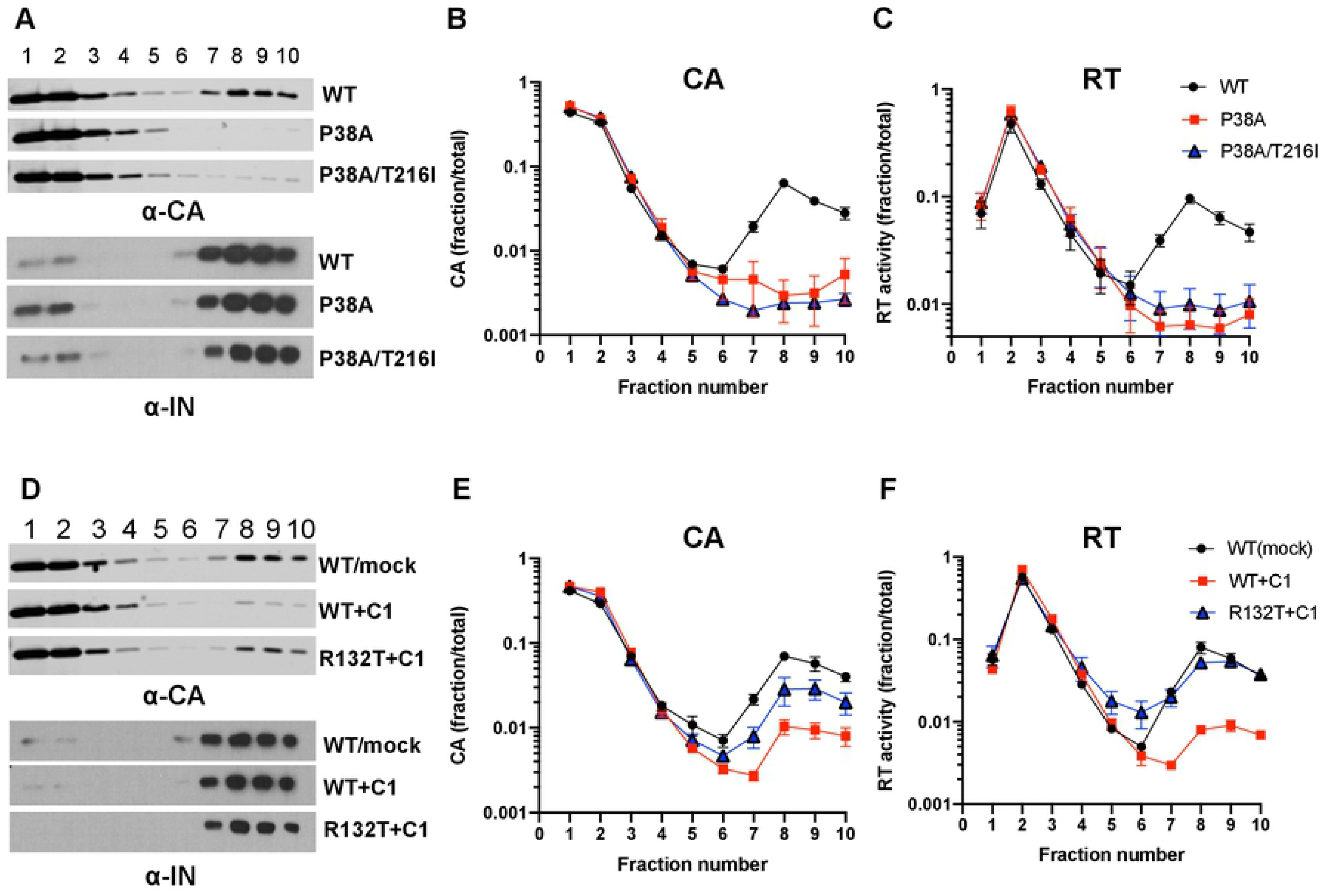
Analyses of compensatory CA mutant viruses. **(A-C)** HIV-1_NL4-3-_derived virions bearing P38A or P38A/T216A substitutions were subjected to equilibrium density sedimentation as in Fig. 4. Ten fractions collected from the top of the sucrose gradients were analyzed for CA (A, B), IN (A) and RT (C) by immunoblotting or a Q-PCR-based RT activity assay. **(D-F)** WT and HIV-1_NL4-3-_derived virions bearing the CA R132T substitution that confers resistance to C1 were generated in the presence of 100 μM C1 and subjected to equilibrium density sedimentation as in Fig. 4. CA (D, E), IN (D) and RT (F) in collected fractions were detected by immunoblotting or a Q-PCR-based RT activity assay. Data in A and D are representative of four independent experiments. Data in B, C, E, F (mean values ± SEM) show the quantitation of CA immunoblots and RT activity over four independent experiments.

### Destabilization of the CA lattice leads to loss of vRNPs in target cells

The impact of CA destabilization on the fates of viral cores in infected cells is poorly studied. Of note, HIV-1 cores can be stabilized by cytosolic extracts in vitro [105], suggesting that core stability in cells may be different from what is predicted from in vitro assays. While elaborate and powerful microscopy-based approaches have recently been employed to fill this gap [21, 22, 24, 25], an important drawback of these approaches is their dependence on indirect labeling of core components and fusion proteins that may impact functionality. In addition, image-based studies have not assessed the impact of CA destabilization on vRNA and IN, which exist at comparatively lower copy numbers in virions. To fill this knowledge gap, we tracked the fates of individual core components in target cells by utilizing our previously developed biochemical assay [95]. In brief, CHO-derived pgsA-745 cells, which lack surface glycosaminoglycans and as a result can be very efficiently infected by VSV-G-pseudotyped viruses, were synchronously infected with WT or CA mutant viruses. Two hours post-infection (hpi), post-nuclear lysates were separated on linear sucrose gradients and collected fractions were analyzed for viral proteins (i.e. CA, IN, RT) and viral genomic RNA or reverse transcription products by immunoblotting or Q-PCR-based assays, respectively.

In cells infected with WT viruses, CA migrated as two populations (Fig. 6A). The first population was present in the top two fractions corresponding to soluble CA proteins that have uncoated from the core or CA that was packaged into virions but was not part of the capsid lattice [26, 106]. A second population of CA was present in fractions 6-8, representing CA that is in complex with vRNPs and RTCs as evident by its co-migration with IN (Fig. 6B), vRNA (Fig. 6C), vDNA (Fig. 6D), and RT (Fig. S3). Notably, in line with the in vitro experiments, CA-destabilizing P38A, K203A, and Q219A substitutions led to loss of CA, whereas the CA-stabilizing E45A substitution yielded similar levels of CA compared to the WT in dense fractions (Fig. 6A). Notably, IN (Fig. 6B) and vRNA (Fig. 6C, S4A) were lost from dense fractions without corresponding increases in soluble fractions upon CA destabilization. These changes expectedly led to substantially lower levels of reverse transcription products accumulating in cells (Fig. 6D, S4B). In line with results of Figures 5A-C, the P38A/T216I substitution did not appreciably restore the amount of CA in dense fractions (Fig. 6A), but did modestly increase the amount of IN (Fig. 6B), vRNA (Fig. 6C, S4A), and vDNA (Fig. 6D, S4B). Core destabilization by C1 similarly reduced the amount of CA in dense fractions (Fig. 6E), which was accompanied by the loss of IN (Fig. 6F) and vRNA (Fig. 6G, S4C). Importantly, the R132T substitution, which confers resistance to C1, largely restored CA, IN, RNA, as well as reverse transcription products in dense fractions for viruses generated in the presence of C1 (Fig. 6E-H, S4C, S4D).

**Figure 6.**
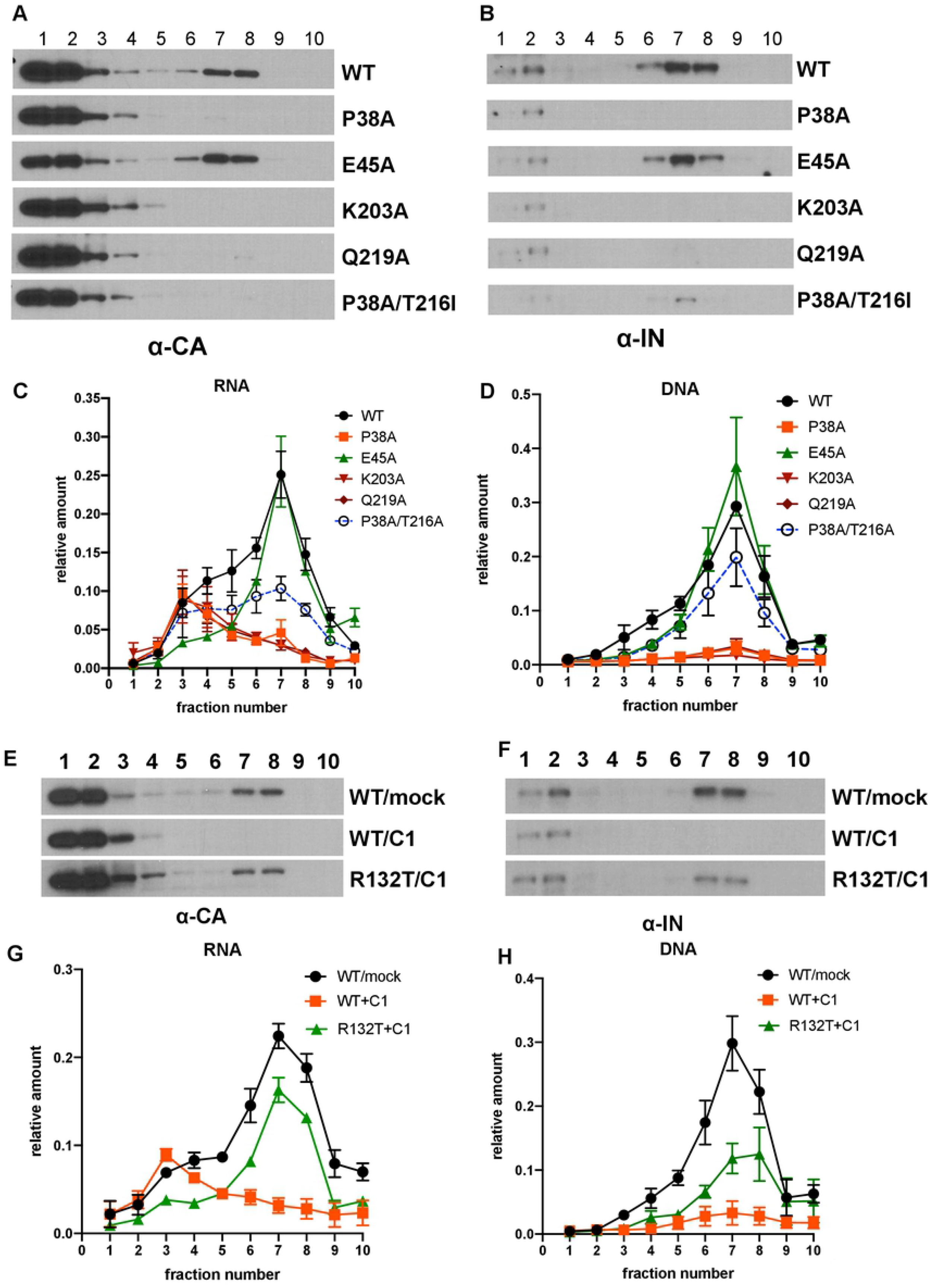
Fates of viral cores with altered stabilities in infected cells. PgsA-745 cells were synchronously infected with VSV-G pseudotyped GFP reporter HIV-1 bearing the indicated CA substitutions (A-D) or generated in the presence of 100 μM C1 (E-H). Cells were processed at 2 hpi as explained in Materials and Methods. Proteins in fractions were analyzed by western blotting using antibodies against CA-p24 (A, E) and IN (B, F). Nucleic acids detected by PCR are shown in panels C and G (vRNA) and D and H (vDNA). Immunoblots in A, B, E, F are representative of three independent experiments. Data in C, D, G, H show mean values from three independent experiments, with error bars representing the SEM.

To test whether accelerated loss of vRNA upon core destabilization holds true in a cell type relevant to HIV-1 infection, MT-4 T-cells were synchronously infected with VSV-G pseudotyped full-length viruses in the presence of the RT inhibitor nevirapine and levels of cell-associated vRNA assessed by Q-RT-PCR. In line with the above findings, premature loss of the genomic vRNA upon CA destabilization was apparent as early as 2 hpi (Fig. 7A). The viral genomic RNA continued to be lost at a faster rate for CA destabilizing mutations and C1 as compared to WT and E45A virus at later times in infection (Fig. 7A). Importantly, equivalent levels of vRNA were recovered from cells when virus entry was blocked through NH_4_Cl treatment (Fig. 7B). These results, together with a previous study that found no difference in the efficiency of VSV-G-mediated viral entry for WT and CA P38A, E45A, and K203A mutant viruses [107], suggest that the observed decrease in vRNA levels is dependent on viral entry. Taken together, these results support our findings from the fate of core assays (Fig. 6) and strongly argue for the role of the CA lattice in protecting vRNPs from degradation in target cells.

**Figure 7.**
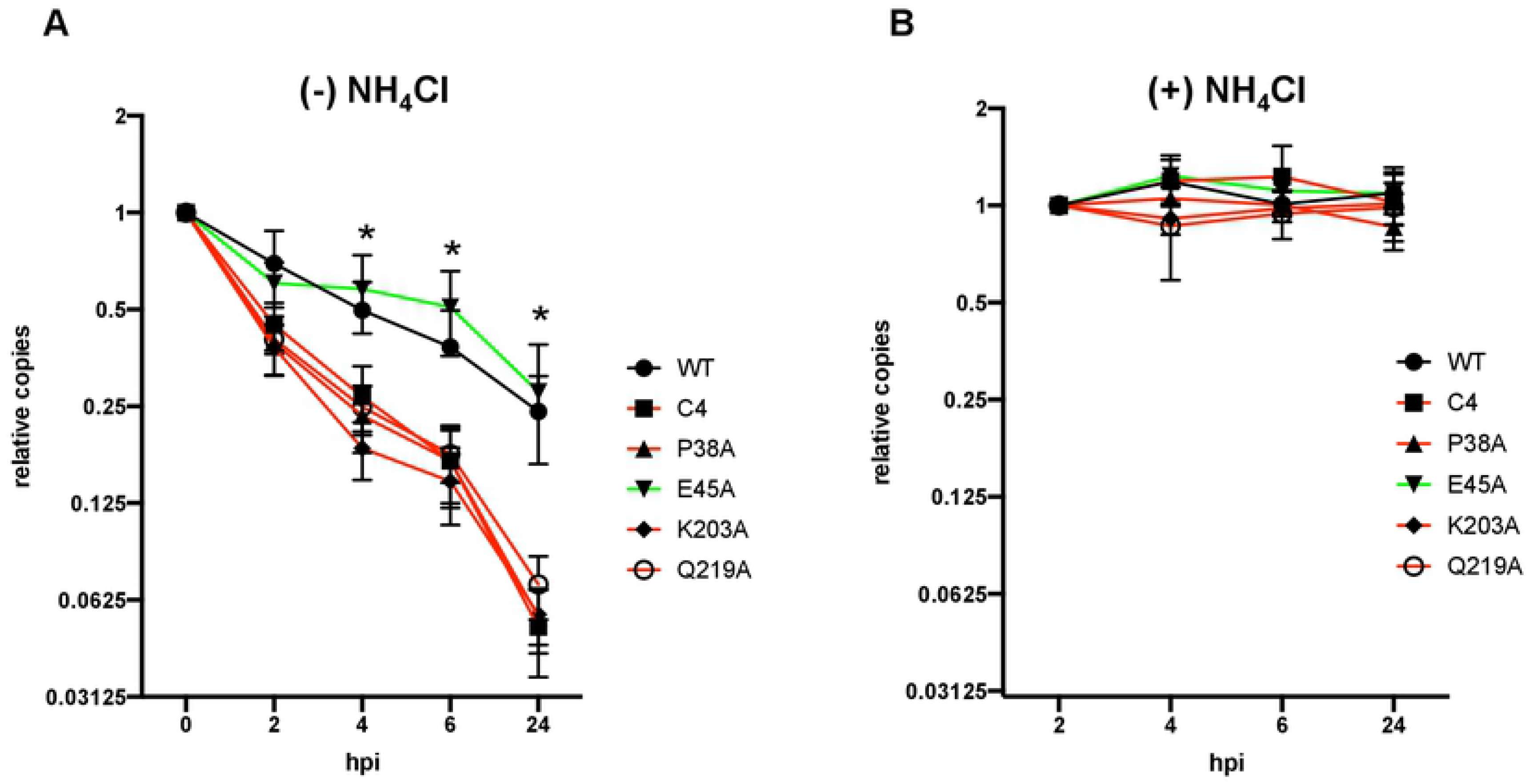
Destabilization of the CA lattice leads to the loss of vRNAs in target cells. **(A,B)** MT4 cells were synchronously infected with HIV-1_NL4-3_/VSV-G at the MOI of 1 i.u./cell for WT (equivalent particle numbers as normalized by RT activity was used for the mutants). Following synchronization, cells were shifted to 37 °C in the presence of 25 μM nevirapine (A) or 25 μM nevirapine and 50 mM ammonium chloride (B). Cell-associated vRNA at indicated time-points was analyzed by Q-RT-PCR. Data in A and B are normalized relative to the amount of RNA detected at T=0 and T=2 hpi, respectively, for each virus. Data are from six independent experiments. Wilcoxon matched-pairs signed rank test was performed to assign significance (*, p<0.05).

### Loss of vRNPs in target cells upon core destabilization is independent of proteasomes

We next tested whether the observed loss of vRNPs is mediated by proteasomes. Of note, it has been previously shown that IN is inherently unstable due to the presence of an N-terminal phenylalanine residue that leads to its proteasomal degradation when expressed alone in cells [108–112]. In agreement with data presented in Fig. 6, CA (Fig. 8A), IN (Fig. 8B), and vRNA (Fig. 8C, S5) were lost from dense fractions for the P38A destabilizing mutant and to a lesser extent for the P38A/T216I mutant virus. Although proteasome inhibition by MG132 treatment during infection modestly increased the levels of these components in mid-gradient fractions (Fig. 8A-C, S5), it had no impact on virus titers (Fig. 8D). Similar results were obtained with K203A and Q219A substitutions whereby CA (Fig. S6A), IN (Fig. S6B), and vRNA (Fig. S6C) levels in mid fractions increased modestly upon proteasome inhibition, yet without detectable changes in virus titers (Fig. S6D).

**Figure 8.**
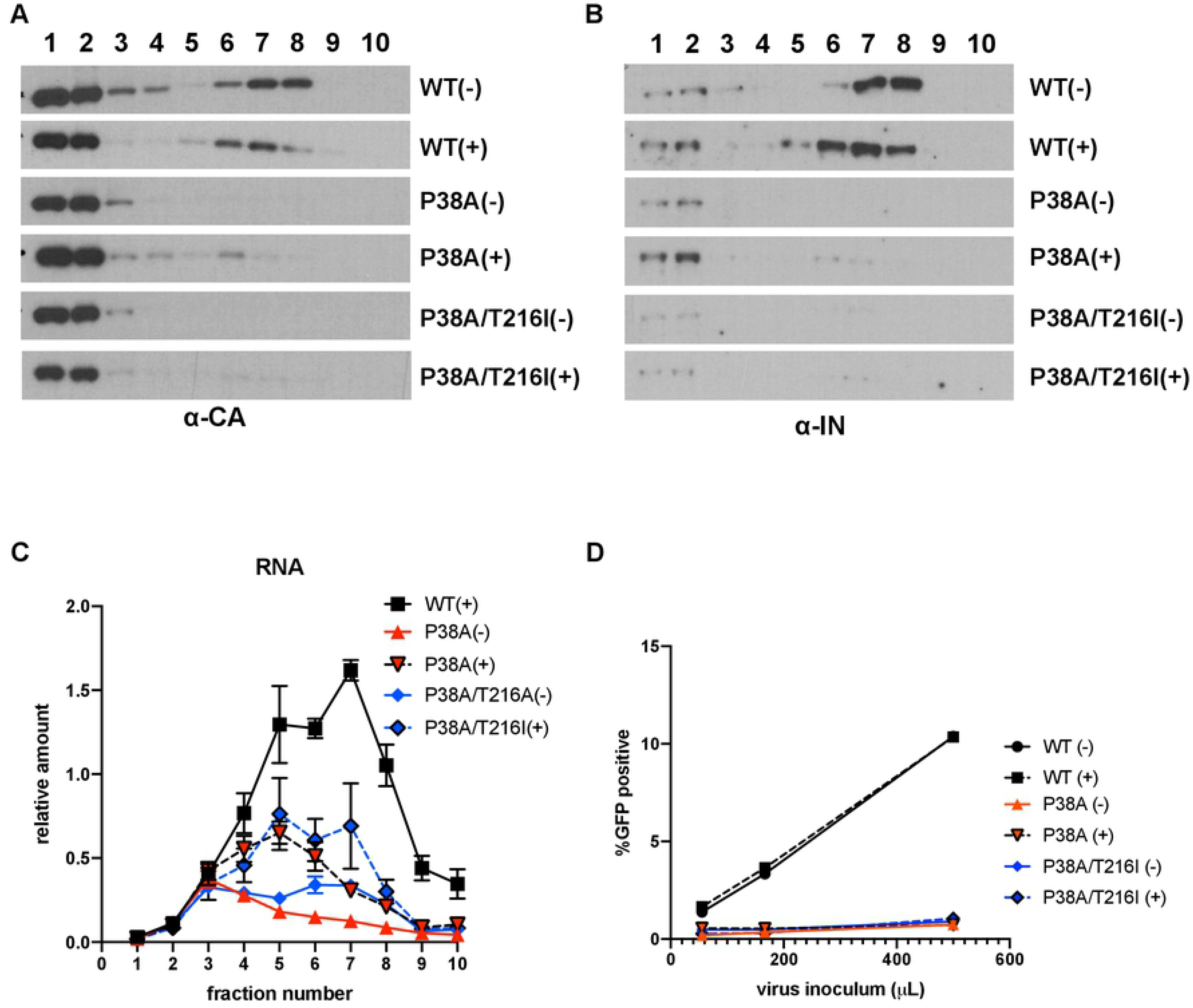
Inhibition of proteasomes does not rescue the degradation of viral cores with decreased stability. PgsA-745 cells were synchronously infected with the indicated WT or CA mutant virus in the absence or presence (− or +) of 2 μM MG132, and infected cells were processed at 2 hpi. Proteins in fractions were analyzed by western blotting using antibodies against CA (A) and IN (B); vRNA was analyzed by Q-RT-PCR (C). Viral titers from a representative experiment are shown in (D). Data are derived from three independent experiments with error bars representing the SEM.

## DISCUSSION

In this study, we utilized complementary approaches to study the impact of CA destabilization on the physical properties of the CA lattice in vitro and in virions, and on the subsequent steps of virus replication in target cells. Our in depth study is the first to causally link how destabilization of the HIV-1 CA lattice leads to reverse transcription defects in target cells. In brief, we found that CA destabilization through multiple mutations and a small molecule compound (C1) all led to faster disassembly of the CA lattice and premature loss, possibly due to degradation, of the vRNA genome and IN in target cells (Fig. 9). Thus, we conclude that protection of vRNPs inside the CA lattice is crucial for reverse transcription as well as subsequent steps in HIV-1 replication.

**Figure 9.**
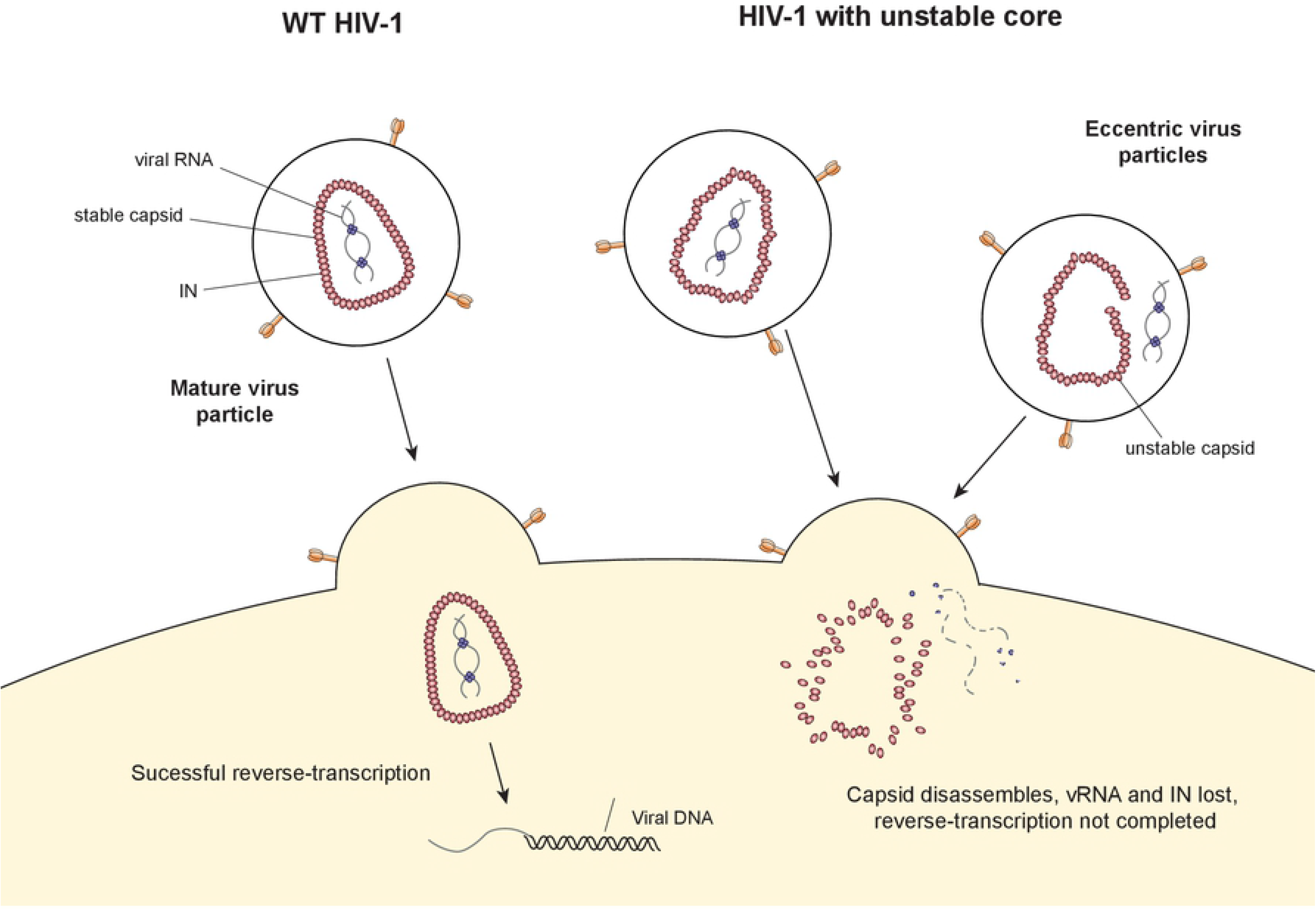
Model of how CA stability affects downstream events in HIV-1 replication.

In terms of the behavior of CA, our findings from fate of core assays are in perfect alignment with previous studies that utilized live cell microscopy approaches [21, 24, 25]. For example, we found that as early as 2 hpi, the majority of virion-associated CA dissociates from the vRNPs of WT viruses. Expectedly, microscopy-based assays that rely on indirect labeling of CA have generally seen a quick loss of CA signal immediately after entry [21, 23–25]. We believe this loss is in part due to uncoating and in part due to the fact that only approximately one-third to one-half of CA monomers in virions form the CA lattice [26, 106], while the remainder diffuses in the cellular milieu upon entry. A third possibility is that CA dissociates from the core during our lysis and fractionation processes. Of note, inclusion of 10 μM IP_6_, which impacts capsid assembly and stability in vitro [10, 11], throughout the fractionation process had no observable impact on the migration behavior of CA and other core components in sucrose gradients (compare Fig. S7 and Fig. 4). Notwithstanding, a small fraction of CA remained associated with vRNPs and the RTC, which was responsive to and was lost upon destabilization of the CA lattice.

Our findings suggest that the main impact of CA destabilizing mutations is on the intrinsic stability of the CA lattice, which is largely in agreement with the core stability assessments in virions and in target cells. One exception was the P38A/T216I double mutant, which we found to be more intrinsically stable than WT in vitro, but was largely unstable in virion- and cell-based stability assays, in agreement with previous findings [101]. Notably, we consistently found higher levels of vRNA, IN, and RT products in dense fractions upon infection of target cells with the P38A/T216I mutant compared to the CA destabilizing P38A mutant, which is consistent with partial rescue of P38A infectivity. Our results suggest that the P38A/T216I substitution may slow down the rate or degree of core disassembly, allowing for intermediate levels of reverse transcription and infection.

Most notably, our study provides the first direct evidence that exposed vRNA and IN are both lost in target cells without the protection of the CA lattice. This is in contrast to a previous study that utilized an IN-superfolder GFP (IN-sfGFP) fusion protein to track RTCs in target cells [113], in which case IN levels did not seem to be affected upon core destabilization [21]. Possible explanations for this discrepancy include the effect of sfGFP fusion on IN function and stability, as well as the artificial introduction of the IN-sfGFP protein into virions through its fusion to Vpr. A separate study observed that the viral genomic RNA labeled with 5-ethynyl uridine was lost quicker from cells upon CA destabilization by the K203A mutant, and curiously, upon CA stabilization by the E45A change [23]. As this study assessed the stability of vRNA in the absence of RT inhibitors, it is possible that the faster loss of vRNAs with the E45A mutant is due to quicker rates of reverse transcription, and hence RNaseH-dependent degradation. Note that we assessed the fates of vRNAs in the presence of RT inhibitor nevirapine to precisely address this problem and circumvent RNaseH-dependent degradation of the vRNA genome during reverse transcription. In addition, we believe that the direct assessment of the behavior of the vRNA genome is another technical strength of our study.

The study of retroviral infection is inherently complicated by the fact that a large fraction of physical particles that enter cells are non-infectious. As a result, it is often assumed that the majority of the infection events studied in biochemical experiments, which depend on the analysis of bulk infected cells, are largely composed of non-infectious viruses [19]. However, we believe that using pgsA-745 cells, which can be very efficiently infected with VSV-G pseudotyped particles (as also observed by others [25]), together with synchronizing the infections, largely mitigates this problem. In fact, while we cannot exclude the possibility that the dense CA containing vRNP complexes that we detect in fate of core assays (Fig, 6, 8) are blocked at downstream events following reverse transcription, they appear to be capable of at least completing reverse transcription. This is based on two observations. First, levels of vRNA detected throughout the gradient decreased substantially if RT inhibitors were omitted during infection, suggesting their efficient reverse transcription (data not shown). This also indicates that the amount of viruses trapped in endosomes, which would appear in middle membrane-containing fractions of the gradients, or viruses being degraded, which would appear in top fractions containing soluble proteins and RNA molecules, is relatively low under these conditions. Second, assuming that the intermediate processing steps work at similar efficiency, the copy numbers of vRNA and vDNA were similar, again suggesting the efficient conversion of vRNA to vDNA by RT.

We have previously shown that HIV-1 IN exhibits a key, non-catalytic role in particle maturation that involves its binding to the vRNA genome [63]. Inhibition of IN-RNA interactions leads to mislocalization of vRNPs outside of the CA lattice [63], and subsequent loss of both the vRNA genome and IN in target cells [90]. Similar loss of the vRNA and IN upon destabilization of the CA lattice, without any apparent effect on IN-RNA and NC-RNA interactions, strongly suggests that it is protection by the CA lattice that matters for the stability of vRNPs as opposed to IN-RNA interaction *per se*.

It remains unknown why the unprotected vRNA and IN are prematurely lost in target cells. One possible hypothesis is that HIV-1 RNAs are inherently unstable due to their AU-rich nucleotide content [114–116], similar to certain cellular mRNAs encoding for cytokines and growth factors [117]. Another is that virion-associated enzymes nick and deadenylate vRNAs in virions [118–120], predisposing them to degradation upon entering target cells. While IN undergoes proteasomal degradation when ectopically expressed alone in cells [108–112], we have found that proteasome inhibition does not rescue the loss of vRNA or IN during infection (Fig. 8 & [90]). Whether the premature loss of unprotected vRNA and IN from infected cells is due to another cellular mechanism or inherent instability of vRNPs remains to be determined.

Our findings may have implications for how HIV-1 nucleic acids are recognized in infected cells by host innate sensors. Shielding of the vRNPs and the resulting reverse transcription products by CA has been proposed to prevent their recognition by cytosolic nucleic acid sensors in immune cells subsets such as dendritic cells and macrophages [53, 56]. For example, perturbation of CA interactions with host cell factors cyclophilin A (CypA) and cleavage and polyadenylation specificity factor subunit 6 (CPSF6) can trigger innate immune responses and interferon (IFN) production in macrophages [55] and monocyte-derived dendritic cells [54]. However the extent of type-I IFN production upon sensing has been variable and dependent on cell-type and culture conditions [121]. For instance, the lack of a robust type I IFN response upon HIV-1 infection of macrophages can be explained by degradation of excess reverse transcription products by the cytosolic exonuclease TREX1 [122], as well as negative regulation of host factors by viral accessory proteins [123]. In other settings, cyclic guanosine monophosphate-adenosine monophosphate synthase (cGAS) and the adaptor protein stimulator of interferon genes (STING), as well as other regulators and downstream effectors, have been proposed to be involved in recognition of HIV-1 DNA [52, 124, 125]. It will be important in the future to determine whether the time window between the exposure of vRNPs and their degradation is sufficiently long to allow innate immune recognition to occur.

Overall, our findings highlight a critical role for the CA lattice in protecting vRNPs from premature degradation in target cells and causally link how CA stability may impact reverse transcription. Given the broad network of essential interactions between CA molecules within the lattice and cellular factors in target cells, HIV-1 CA is emerging as a viable new target for anti-retroviral therapy [17]. Compounds that target CA can disrupt the assembly of the CA lattice and particle morphogenesis [91, 126–129], alter the stability of the CA lattice and/or uncoating [91, 130–132], and inhibit reverse transcription [91, 126, 129, 130, 132, 133] and nuclear entry [134–136] in target cells. Expectedly, CA is highly sensitive to mutations [137], making it an exceptionally viable drug target as resistance mutations would likely come at a high fitness cost to the virus.

## MATERIALS AND METHODS

### Chemicals and reagents

Standard laboratory chemicals were obtained from reputable suppliers such as Sigma-Aldrich. The RT inhibitor nevirapine was obtained from the NIH AIDS Repository, while compound C1 was synthesized as described previously [90].

### Plasmids

The pNLGP plasmid consisting of the HIV-1_NL4-3_ -derived Gag-Pol sequence inserted into the pCR/V1 plasmid backbone [138] and the CCGW vector genome plasmid carrying a GFP reporter under the control of the CMV promoter [139, 140] were previously described. Mutations in the CA coding sequence were introduced into both the pNLGP plasmid and pNL4-3 by overlap extension PCR. Briefly, forward and reverse primers containing CA mutations were used in PCR reactions with antisense and sense outer primers containing unique restriction endonuclease sites (EcoRI-sense, NotI-antisense for NLGP and BssHII-sense-SphI-antisense or SphI-sense-AgeI-antisense for pNL4-3), respectively. The resulting fragments containing CA mutations were mixed at 1:1 ratio and overlapped subsequently using the outer sense and antisense primer pairs. PCR products were digested with the corresponding restriction endonucleases and ligated with appropriately digested pNLGP or pNL4-3 plasmid vector fragments. Presence of engineered mutations and lack of unwanted extraneous mutations were verified by Sanger sequencing.

### Cells and viruses

HEK293T cells (ATCC CRL-11268) and HeLa-derived TZM-bl cells (NIH AIDS Reagent Program) were maintained in Dulbecco’s modified Eagle’s medium supplemented with 10% fetal bovine serum. CHO K1-derived pgsA-745 cells (CRL-2242, ATCC) were maintained in Dulbecco’s modified Eagle’s / F12 (1:1) media supplemented with 10% fetal bovine serum and 1 mM L-glutamine. MT-4 T cells (NIH AIDS Reagents) were grown in RPMI media supplemented with 10% fetal bovine serum. Vesicular stomatitis virus G protein (VSV-G)-pseudotyped virus-like particles (VLPs) were produced by transfection of HEK293T cells with pNLGP-derived plasmids, the CCGW vector genome carrying GFP, and VSV-G expression plasmid at a ratio of 5:5:1, respectively, using polyethyleneimine (PolySciences, Warrington, PA). VSV-G-pseudotyped viruses were produced by transfecting HEK293T cells with the pNL4-3-derived plasmids and VSV-G plasmid at a ratio of 4:1 (pNL4-3:VSV-G).

### Immunoblotting

Viral and cell lysates were resuspended in SDS sample buffer and separated by electrophoresis on Bolt 4-12% Bis-Tris Plus gels (Life Technologies), blotted onto nitrocellulose membranes, and probed with the following antibodies: mouse monoclonal anti-HIV p24 antibody (183-H12-5C, NIH AIDS reagents), mouse monoclonal anti-HIV IN antibody [141], rabbit polyclonal anti-HIV IN antibody raised in-house against the Q44-LKGEAMHGQVD-C56 peptide. Blots were then probed with fluorophore-conjugated secondary antibodies (LI-COR) and scanned using a LI-COR Odyssey system. IN and CA levels in virions were quantified using the Image Studio software.

### Equilibrium density sedimentation of virion core components *in vitro*

Equilibrium density sedimentation of virion core components was performed as previously described [90]. Briefly, HEK293T cells grown on 10-cm dishes were transfected with NLGP or derivative plasmids. Two days post-transfection, cell-free virions collected from cell culture supernatants were pelleted through a 20% sucrose cushion. Pelleted VLPs were resuspended in 1x PBS and treated with 0.5% Triton X-100 for 2 min at room temperature. Immediately after, samples were layered on top of 30-70% linear sucrose gradients in 1X STE buffer (100 mM NaCl, 10 mM Tris-Cl (pH 8.0), 1 mM EDTA) and centrifuged for 16 h at 4°C and 28500 rpm, respectively, using an SW55Ti rotor. Fractions (500 μL) collected from the top were analyzed for IN by immunoblotting using a mouse monoclonal anti-IN antibody [141], anti-HIV p24 antibody (183-H12-5C, NIH AIDS reagents), rabbit polyclonal anti-MA antibody (4811, NIH AIDS reagents), and Q-PCR-based assays for RT activity [142] and vRNA.

### Analysis of virion core components in infected cells

Biochemical analysis of retroviral cores in infected cells was performed as described previously [95]. Briefly, pgsA-745 cells were mixed with VSV-G pseudotyped single cycle GFP-reporter viruses or its derivatives at 4°C. Following the removal of virus inoculum and extensive washes with 1X PBS, cells were incubated at 37°C for 2 h. For analysis of vRNA, 25 μM nevirapine was included throughout the infections to prevent its degradation during reverse transcription due to RNase H activity. Post-nuclear supernatants were separated by ultracentrifugation on 10-50% linear sucrose gradients using a SW50.1 rotor at 30,000 rpm for 1 h. Ten 500 μl fractions from the top of the gradient were collected, and CA, IN, RT activity, vRNA and vDNA in each fraction were analyzed by either immunoblotting or Q-PCR as above and detailed in [95].

### Virus production and transmission electron microscopy

Cell-free HIV-1 virions were isolated from transfected HEK293T cells. Briefly, cells grown in two 15-cm dishes (10^7^ cells per dish) were transfected with 30 μg full-length proviral plasmid DNA containing the WT sequence or indicated CA mutations using PolyJet DNA transfection reagent as recommended by the manufacturer (SignaGen Laboratories). Two days after transfection, cell supernatants were filtered through 0.22 μm filters and pelleted by ultracentrifugation using a Beckman SW32-Ti rotor at 26,000 rpm for 2 h at 4 °C. Fixative (2.5% glutaraldehyde, 1.25% paraformaldehyde, 0.03% picric acid, 0.1 M sodium cacodylate, pH 7.4) was gently added to resulting pellets, and samples were incubated overnight at 4 °C. The following steps were conducted at the Harvard Medical School Electron Microscopy core facility. Samples were washed with 0.1 M sodium cacodylate, pH 7.4 and postfixed with 1% osmium tetroxide /1.5% potassium ferrocyanide for 1 h, washed twice with water, once with maleate buffer (MB), and incubated in 1% uranyl acetate in MB for 1 h. Samples washed twice with water were dehydrated in ethanol by subsequent 10 min incubations with 50%, 70%, 90%, and then twice with 100%. The samples were then placed in propyleneoxide for 1 h and infiltrated overnight in a 1:1 mixture of propyleneoxide and TAAB Epon (Marivac Canada Inc.). The following day the samples were embedded in TAAB Epon and polymerized at 60 °C for 48 h. Ultrathin sections (about 60 nm) were cut on a Reichert Ultracut-S microtome, transferred to copper grids stained with lead citrate, and examined in a JEOL 1200EX transmission electron microscope with images recorded on an AMT 2k CCD camera. Images were captured at 30,000 x magnification, and over 100 viral particles per sample were counted by visual inspection.

### Nano differential scanning fluorimetry (NanoDSF) analysis of CA assemblies

Purified HIV-1 CA proteins (WT, P38A, P38A/T216I, E45A, K203A, and Q219A) were obtained using published protocols [143]. CA tubes were assembled by incubating protein (~10 mg/mL) in 50 mM Tris, pH 8.0, 1 M NaCl, 20 mM β-mercaptoethanol for 2 h at 37 °C. Unassembled protein were removed by centrifugation, and samples were then diluted 10-fold into the same buffer and incubated for 10 min at room temperature prior to loading onto nano-capillaries. NanoDSF profiles were measured with a Tycho system (Nanotemper). The first derivative profiles were then fit as a sum of Gaussian curves in Excel (Microsoft).

### Analysis of vRNA in synchronously infected MT4 cells

MT4 cells (3-6×10^6^) were cooled to 4°C and infected with HIV-1_NL4-3_/VSV-G in the presence of 5 μM polybrene and 25 μM nevirapine. Equivalent number of particles for CA mutant viruses (as normalized by RT activity) was used to infect cells in parallel. Cells were incubated with viruses at 4°C for 30 min to allow binding, followed by three washes with ice-cold 1xPBS to remove unbound virus. Cells were then shifted to 37°C in the presence of 25 μM nevirapine to allow virus entry. In some experiments 50 mM ammonium chloride was included at this stage to prevent endosome acidification, and hence viral entry. Infected cells were collected at 0, 2, 4, 6 and 24 hpi and RNA was extracted by Trizol. Resulting RNA was reverse transcribed and subjected to Q-PCR analysis for viral genomic RNA.

---

**Figure S1-Supplementary to Figure 4:**
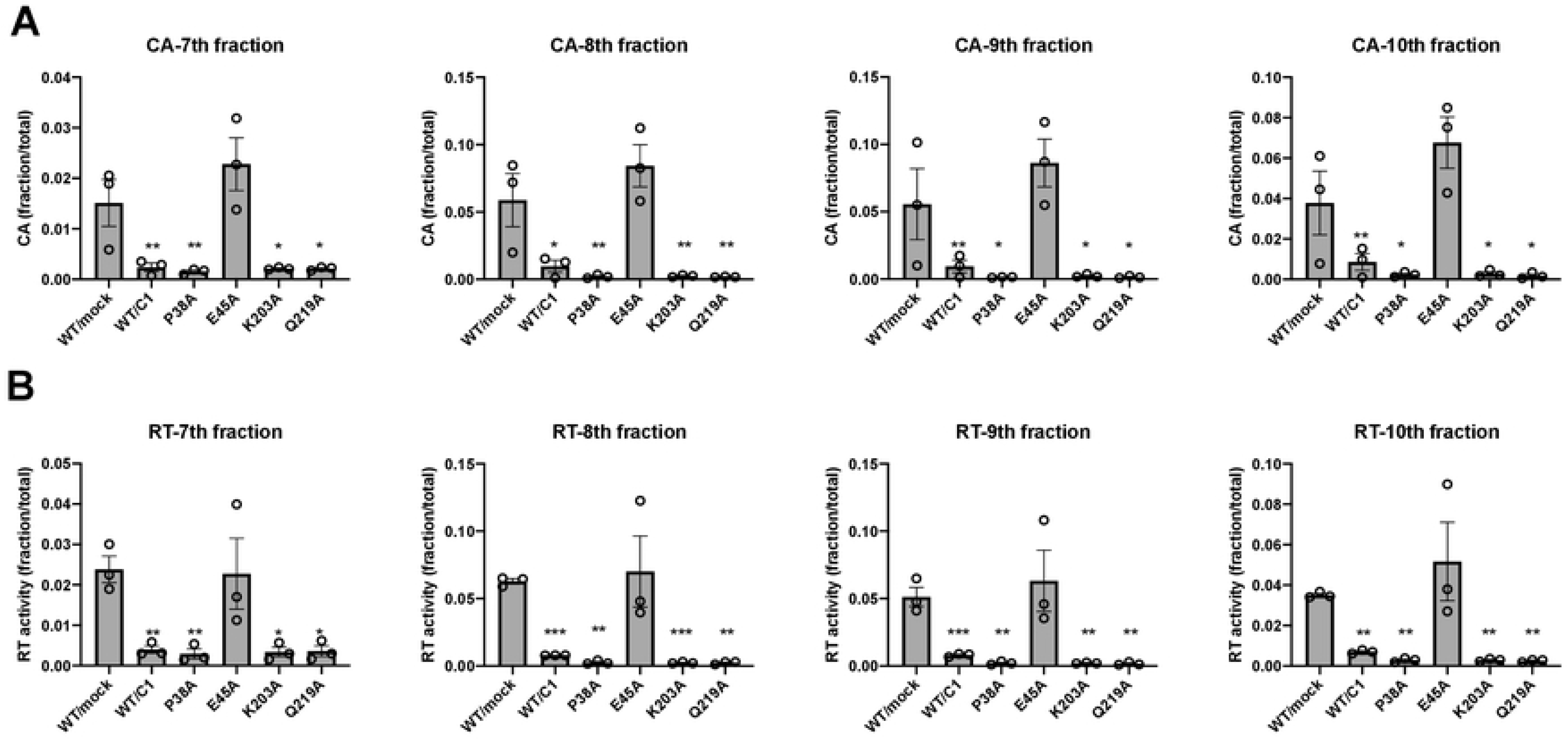
Statistical analysis of the data presented in Figure 4. Ratio paired one-tailed t-test was performed to determine statistically significant decreases (black asterisks) compared to WT/mock sample in CA (A) and RT (B) levels in fractions 7-10. p<0.05 (*), p<0.01 (**), p<0.001 (***).

**Figure S2-Supplementary to Figure 5:**
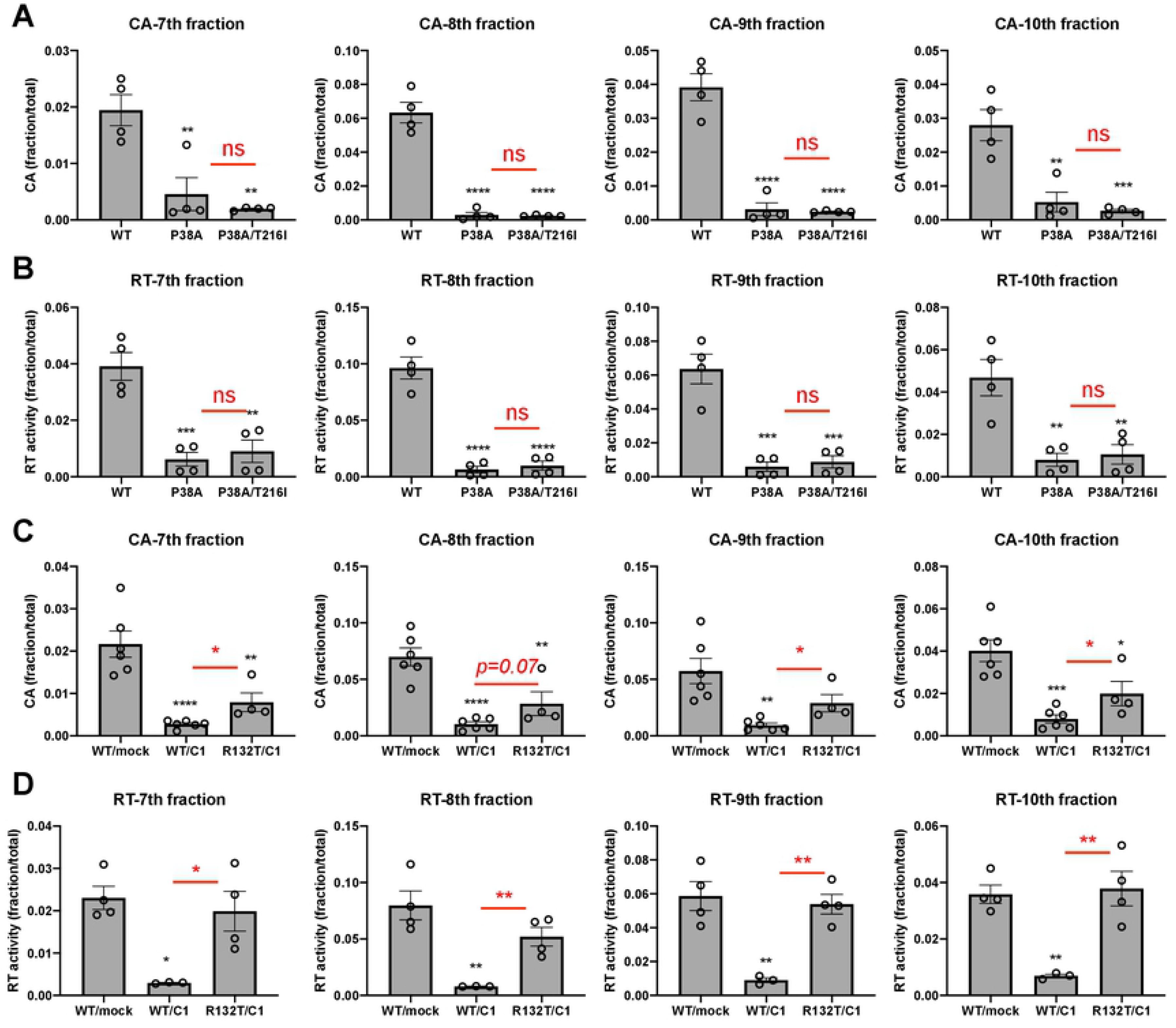
Statistical analysis of the data presented in Figure 5. Ordinary one-way ANOVA analysis was performed to determine statistically significant changes in CA (A, C) and RT (B, D) levels in fractions 7-10. Significant differences compared to the WT sample are denoted in black asterisks. Differences between P38A vs. P38A/T216I mutants (A, B), and WT/C1 vs. R132T/C1 (C, D) samples, were assessed by unpaired two-tailed t-test. These significance levels are indicated in red asterisks. ns, not significant (p>0.05); p<0.05 (*), p<0.01 (**), p<0.001 (***), p<0.0001(****).

**Figure S3-Supplementary to Figure 6:**
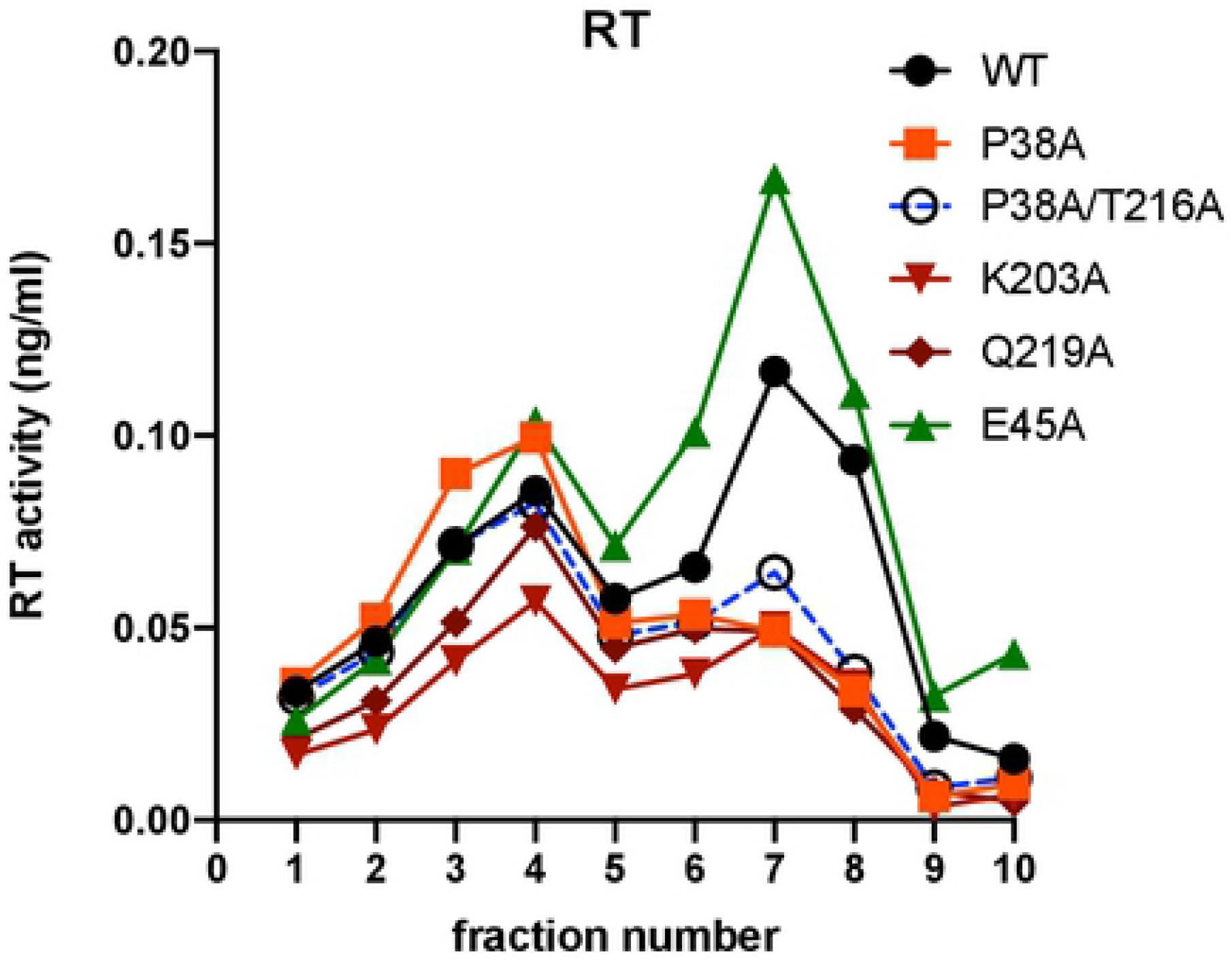
Fates of viral RT with altered stabilities in infected cells. PgsA-745 cells were synchronously infected with VSV-G pseudotyped GFP reporter HIV-1, bearing the indicated CA substitutions. RT activity in fractions were analyzed by a Q-PCR-based assay. Data is from three independent experiments.

**Figure S4-Supplementary to Figure 6:**
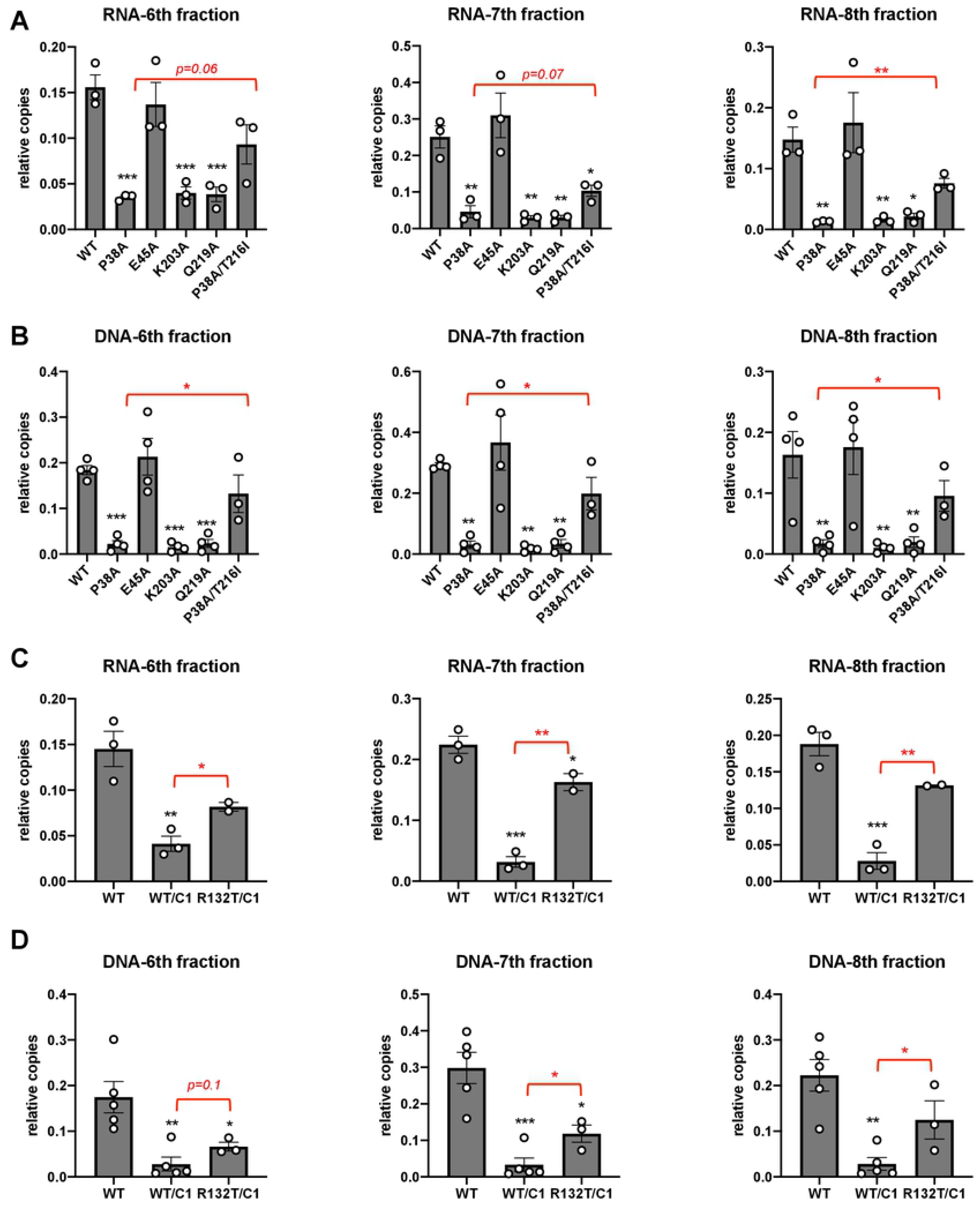
Statistical analysis of the data presented in Figure 6. Ordinary one-way ANOVA analysis was performed to determine statistically significant changes in RNA (A, C) and DNA levels (B, D) in fractions 6-8. Significant differences compared to the WT sample are denoted in black asterisks. Differences between P38A vs. P38A/T216I mutants (A, B) and WT/C1 and R132T/C1 (C, D) samples were assessed by unpaired two-tailed t-test, with these significance levels indicated in red. p<0.05 (*), p<0.01 (**), p<0.001 (***).

**Figure S5-Supplementary to Figure 8:**
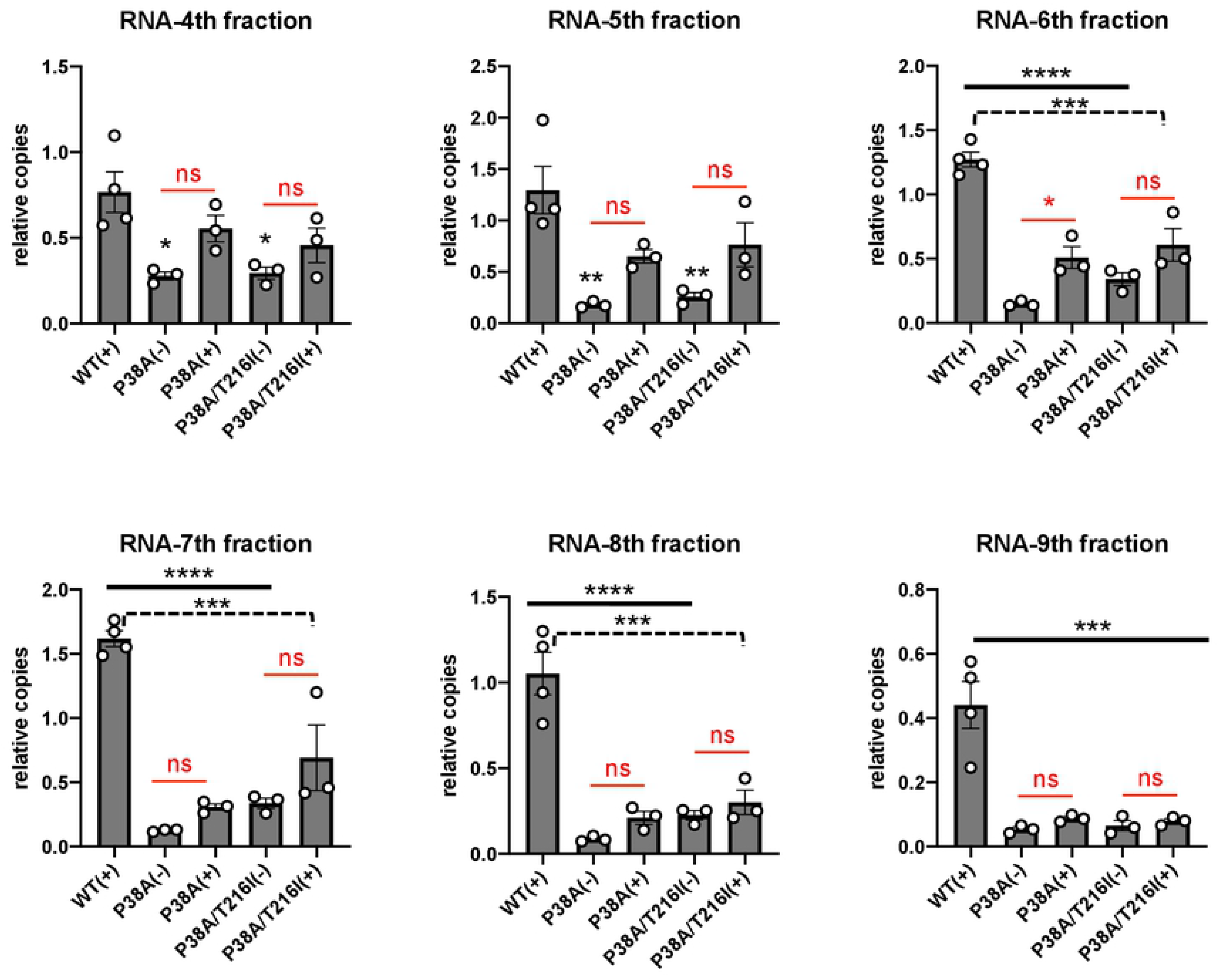
Statistical analysis of the data presented in Figure 8. Ordinary one-way ANOVA analysis was performed to determine statistically significant changes in RNA levels in fractions 4-9. Differences relative to the WT (+) sample are denoted in black asterisks, and differences resulting from MG132 treatment are indicated in red. ns, not significant (p>0.05); p<0.05 (*), p<0.01 (**), p<0.001 (***), p<0.0001(****).

**Figure S6-Supplementary to Figure 8.**
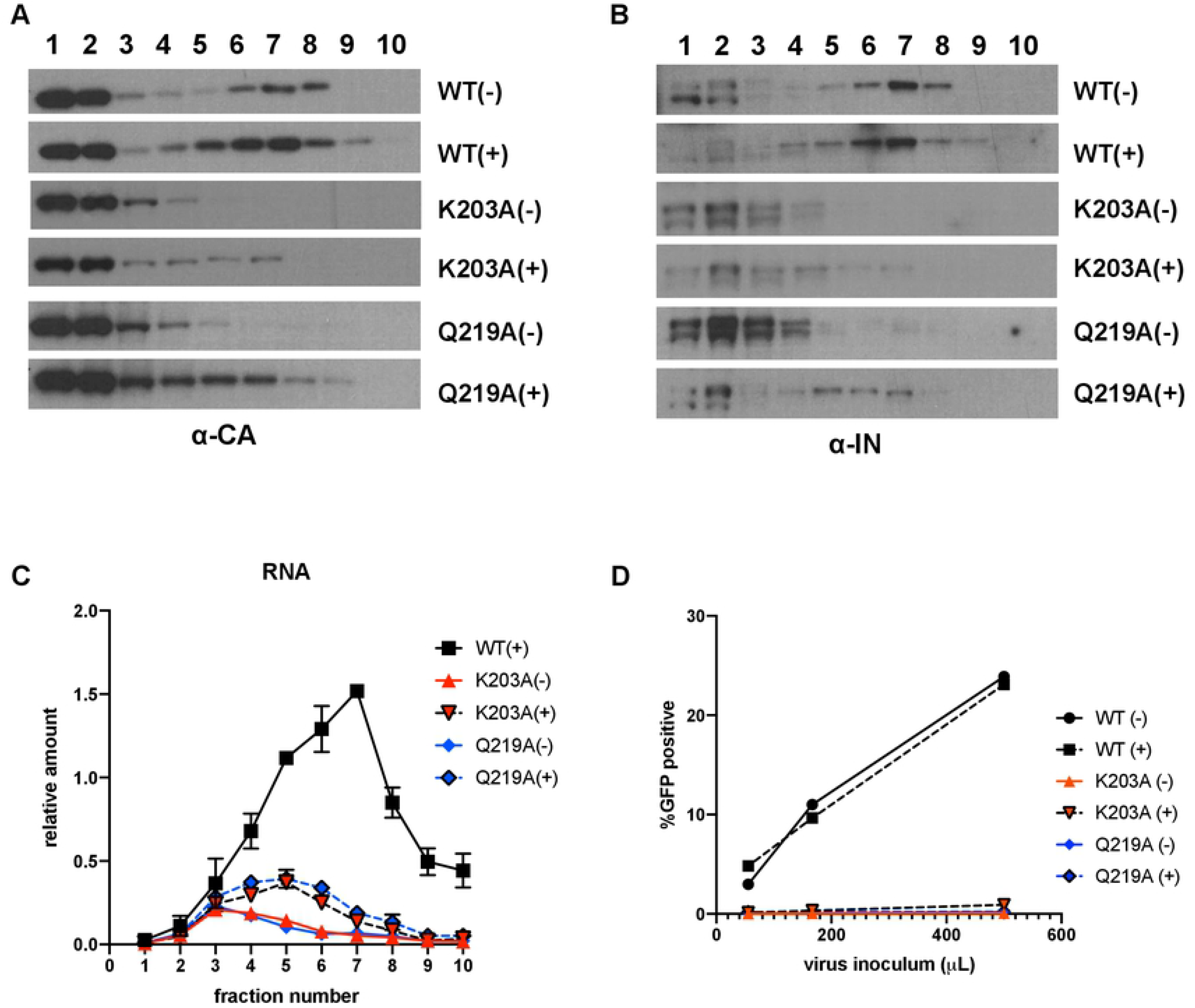
Inhibition of proteasomes does not rescue the degradation of viral cores with decreased stability. PgsA-745 cells were synchronously infected with VSV-G pseudotyped GFP reporter HIV-1 bearing the indicated CA substitutions in the absence or presence of 2 μM MG132, and infected cells were processed at 2 hpi. Proteins in fractions were analyzed by western blotting using antibodies against CA (A) and IN (B). Viral RNA in collected fractions was analyzed by Q-RT-PCR (C). Viral titers from a representative experiments are shown in (D). Data are derived from three independent experiments with error bars representing the SEM.

**Figure S7.**
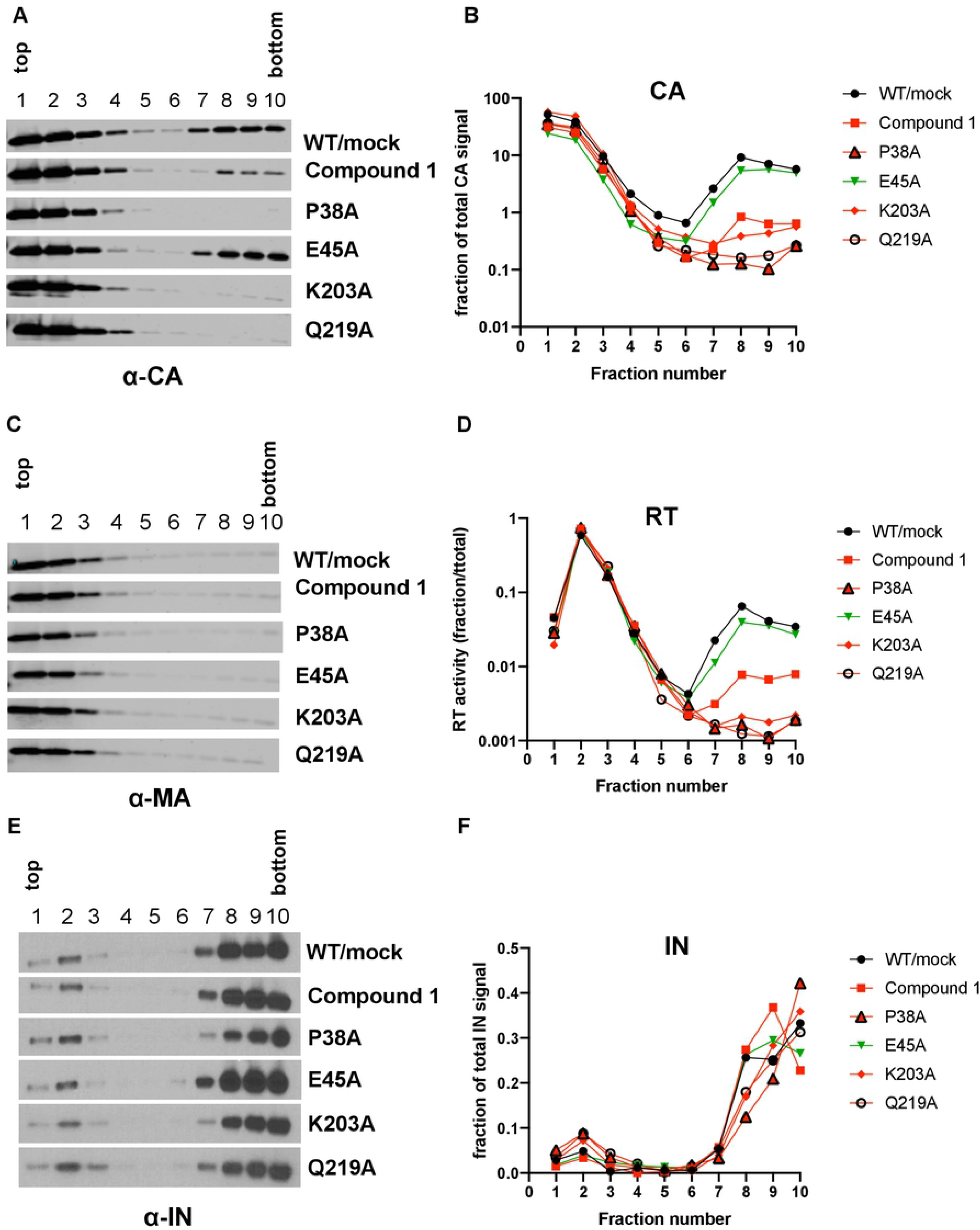
Effects of CA mutations and C1 on the stability of HIV-1 cores in vitro and the impact of IP_6_. Indicated CA mutant viruses or WT grown in the presence of 100 μM C1 were subjected to equilibrium density sedimentation following treatment with 0.5% Triton. Both the lysis buffer and gradients were supplemented with 10 mM IP_6_. Ten fractions collected from the top of the sucrose gradients were analyzed for CA (A, B), MA (C), RT (D), and IN (E, F) by immunoblotting or a Q-PCR-based RT activity assay. Data are representative of two independent experiments.

## ACKNOWLEDGEMENTS

We thank Dr. Michael Malim for providing the anti-IN monoclonal antibody. This study was supported by NIH grants U54AI150470 (the Center for HIV RNA Studies) and AI150497 to SBK, F31AI143389 fellowship to JLE, AI129678 and AI150479 to OP, and P50AI150481 (Pittsburgh Center for HIV Protein Interactions) and AI070042 to ANE.

## REFERENCES

1. Freed, E.O., HIV-1 assembly, release and maturation. Nat Rev Microbiol, 2015. 13(8): p. 484–96.

2. Sundquist, W.I. and H.G. Krausslich, HIV-1 assembly, budding, and maturation. Cold Spring Harb Perspect Med, 2012. 2(7): p. a006924.

3. Mattei, S., F.K. Schur, and J.A. Briggs, Retrovirus maturation-an extraordinary structural transformation. Curr Opin Virol, 2016. 18: p. 27–35.

4. Pornillos, O., et al., X-ray structures of the hexameric building block of the HIV capsid. Cell, 2009. 137(7): p. 1282–92.

5. Pornillos, O., B.K. Ganser-Pornillos, and M. Yeager, Atomic-level modelling of the HIV capsid. Nature, 2011. 469(7330): p. 424–7.

6. Briggs, J.A. and H.G. Krausslich, The molecular architecture of HIV. J Mol Biol, 2011. 410(4): p. 491–500.

7. Ganser-Pornillos, B.K., M. Yeager, and O. Pornillos, Assembly and architecture of HIV. Adv Exp Med Biol, 2012. 726: p. 441–65.

8. Perilla, J.R. and A.M. Gronenborn, Molecular Architecture of the Retroviral Capsid. Trends Biochem Sci, 2016. 41(5): p. 410–420.

9. Mattei, S., et al., The structure and flexibility of conical HIV-1 capsids determined within intact virions. Science, 2016. 354(6318): p. 1434–1437.

10. Dick, R.A., et al., Inositol phosphates are assembly co-factors for HIV-1. Nature, 2018. 560(7719): p. 509–512.

11. Mallery, D.L., et al., IP6 is an HIV pocket factor that prevents capsid collapse and promotes DNA synthesis. Elife, 2018. 7.

12. Byeon, I.J., et al., Structural convergence between Cryo-EM and NMR reveals intersubunit interactions critical for HIV-1 capsid function. Cell, 2009. 139(4): p. 780–90.

13. Forshey, B.M., et al., Formation of a human immunodeficiency virus type 1 core of optimal stability is crucial for viral replication. J Virol, 2002. 76(11): p. 5667–77.

14. Yufenyuy, E.L. and C. Aiken, The NTD-CTD intersubunit interface plays a critical role in assembly and stabilization of the HIV-1 capsid. Retrovirology, 2013. 10: p. 29.

15. Zhao, G., et al., Mature HIV-1 capsid structure by cryo-electron microscopy and all-atom molecular dynamics. Nature, 2013. 497(7451): p. 643–6.

16. von Schwedler, U.K., et al., Functional surfaces of the human immunodeficiency virus type 1 capsid protein. J Virol, 2003. 77(9): p. 5439–50.

17. Carnes, S.K., J.H. Sheehan, and C. Aiken, Inhibitors of the HIV-1 capsid, a target of opportunity. Curr Opin HIV AIDS, 2018. 13(4): p. 359–365.

18. Arhel, N., Revisiting HIV-1 uncoating. Retrovirology, 2010. 7: p. 96.

19. Campbell, E.M. and T.J. Hope, HIV-1 capsid: the multifaceted key player in HIV-1 infection. Nat Rev Microbiol, 2015. 13(8): p. 471–83.

20. Ambrose, Z. and C. Aiken, HIV-1 uncoating: connection to nuclear entry and regulation by host proteins. Virology, 2014. 454-455: p. 371–9.

21. Francis, A.C., et al., Time-Resolved Imaging of Single HIV-1 Uncoating In Vitro and in Living Cells. PLoS Pathog, 2016. 12(6): p. e1005709.

22. Mamede, J.I. and T.J. Hope, Detection and Tracking of Dual-Labeled HIV Particles Using Wide-Field Live Cell Imaging to Follow Viral Core Integrity. Methods Mol Biol, 2016. 1354: p. 49–59.

23. Xu, H., et al., Evidence for biphasic uncoating during HIV-1 infection from a novel imaging assay. Retrovirology, 2013. 10: p. 70.

24. Francis, A.C. and G.B. Melikyan, Single HIV-1 Imaging Reveals Progression of Infection through CA-Dependent Steps of Docking at the Nuclear Pore, Uncoating, and Nuclear Transport. Cell Host Microbe, 2018. 23(4): p. 536–548 e6.

25. Mamede, J.I., et al., Early cytoplasmic uncoating is associated with infectivity of HIV-1. Proc Natl Acad Sci U S A, 2017. 114(34): p. E7169–E7178.

26. Briggs, J.A., et al., The stoichiometry of Gag protein in HIV-1. Nat Struct Mol Biol, 2004. 11(7): p. 672–5.

27. Briggs, J.A., et al., Structural organization of authentic, mature HIV-1 virions and cores. EMBO J, 2003. 22(7): p. 1707–15.

28. Lanman, J., et al., Key interactions in HIV-1 maturation identified by hydrogen-deuterium exchange. Nat Struct Mol Biol, 2004. 11(7): p. 676–7.

29. Marquez, C.L., et al., Kinetics of HIV-1 capsid uncoating revealed by single-molecule analysis. Elife, 2018. 7.

30. Hulme, A.E. and T.J. Hope, The cyclosporin A washout assay to detect HIV-1 uncoating in infected cells. Methods Mol Biol, 2014. 1087: p. 37–46.

31. Hulme, A.E., O. Perez, and T.J. Hope, Complementary assays reveal a relationship between HIV-1 uncoating and reverse transcription. Proc Natl Acad Sci U S A, 2011. 108(24): p. 9975–80.

32. Arfi, V., et al., Characterization of the behavior of functional viral genomes during the early steps of human immunodeficiency virus type 1 infection. J Virol, 2009. 83(15): p. 7524–35.

33. Yamashita, M. and A.N. Engelman, Capsid-Dependent Host Factors in HIV-1 Infection. Trends Microbiol, 2017. 25(9): p. 741–755.

34. Hilditch, L. and G.J. Towers, A model for cofactor use during HIV-1 reverse transcription and nuclear entry. Curr Opin Virol, 2014. 4: p. 32–6.

35. Lee, K., et al., Flexible use of nuclear import pathways by HIV-1. Cell Host Microbe, 2010. 7(3): p. 221–33.

36. Matreyek, K.A. and A. Engelman, The requirement for nucleoporin NUP153 during human immunodeficiency virus type 1 infection is determined by the viral capsid. J Virol, 2011. 85(15): p. 7818–27.

37. Matreyek, K.A. and A. Engelman, Viral and cellular requirements for the nuclear entry of retroviral preintegration nucleoprotein complexes. Viruses, 2013. 5(10): p. 2483–511.

38. Matreyek, K.A., et al., Nucleoporin NUP153 phenylalanine-glycine motifs engage a common binding pocket within the HIV-1 capsid protein to mediate lentiviral infectivity. PLoS Pathog, 2013. 9(10): p. e1003693.

39. Schaller, T., et al., HIV-1 capsid-cyclophilin interactions determine nuclear import pathway, integration targeting and replication efficiency. PLoS Pathog, 2011. 7(12): p. e1002439.

40. Stultz, R.D., J.J. Cenker, and D. McDonald, Imaging HIV-1 Genomic DNA from Entry through Productive Infection. J Virol, 2017. 91(9).

41. Yamashita, M. and M. Emerman, Capsid is a dominant determinant of retrovirus infectivity in nondividing cells. J Virol, 2004. 78(11): p. 5670–8.

42. Burdick, R.C., et al., Dynamics and regulation of nuclear import and nuclear movements of HIV-1 complexes. PLoS Pathog, 2017. 13(8): p. e1006570.

43. Chen, N.Y., et al., HIV-1 capsid is involved in post-nuclear entry steps. Retrovirology, 2016. 13: p. 28.

44. Chin, C.R., et al., Direct Visualization of HIV-1 Replication Intermediates Shows that Capsid and CPSF6 Modulate HIV-1 Intra-nuclear Invasion and Integration. Cell Rep, 2015. 13(8): p. 1717–31.

45. Hulme, A.E., et al., Complementary Assays Reveal a Low Level of CA Associated with Viral Complexes in the Nuclei of HIV-1-Infected Cells. J Virol, 2015. 89(10): p. 5350–61.

46. Peng, K., et al., Quantitative microscopy of functional HIV post-entry complexes reveals association of replication with the viral capsid. Elife, 2014. 3: p. e04114.

47. Burdick, R.C., et al., HIV-1 uncoats in the nucleus near sites of integration. Proc Natl Acad Sci U S A, 2020. 117(10): p. 5486–5493.

48. Koh, Y., et al., Differential effects of human immunodeficiency virus type 1 capsid and cellular factors nucleoporin 153 and LEDGF/p75 on the efficiency and specificity of viral DNA integration. J Virol, 2013. 87(1): p. 648–58.

49. Ocwieja, K.E., et al., HIV integration targeting: a pathway involving Transportin-3 and the nuclear pore protein RanBP2. PLoS Pathog, 2011. 7(3): p. e1001313.

50. Serrao, E. and A.N. Engelman, Sites of retroviral DNA integration: From basic research to clinical applications. Crit Rev Biochem Mol Biol, 2016. 51(1): p. 26–42.

51. Sowd, G.A., et al., A critical role for alternative polyadenylation factor CPSF6 in targeting HIV-1 integration to transcriptionally active chromatin. Proc Natl Acad Sci U S A, 2016. 113(8): p. E1054–63.

52. Gao, D., et al., Cyclic GMP-AMP synthase is an innate immune sensor of HIV and other retroviruses. Science, 2013. 341(6148): p. 903–6.

53. Lahaye, X. and N. Manel, Viral and cellular mechanisms of the innate immune sensing of HIV. Curr Opin Virol, 2015. 11: p. 55–62.

54. Lahaye, X., et al., The capsids of HIV-1 and HIV-2 determine immune detection of the viral cDNA by the innate sensor cGAS in dendritic cells. Immunity, 2013. 39(6): p. 1132–42.

55. Rasaiyaah, J., et al., HIV-1 evades innate immune recognition through specific cofactor recruitment. Nature, 2013. 503(7476): p. 402–405.

56. Towers, G.J. and M. Noursadeghi, Interactions between HIV-1 and the cell-autonomous innate immune system. Cell Host Microbe, 2014. 16(1): p. 10–8.

57. Fitzon, T., et al., Proline residues in the HIV-1 NH2-terminal capsid domain: structure determinants for proper core assembly and subsequent steps of early replication. Virology, 2000. 268(2): p. 294–307.

58. Reicin, A.S., et al., The role of Gag in human immunodeficiency virus type 1 virion morphogenesis and early steps of the viral life cycle. J Virol, 1996. 70(12): p. 8645–52.

59. Tang, S., et al., Human immunodeficiency virus type 1 N-terminal capsid mutants that exhibit aberrant core morphology and are blocked in initiation of reverse transcription in infected cells. J Virol, 2001. 75(19): p. 9357–66.

60. Cosnefroy, O., P.J. Murray, and K.N. Bishop, HIV-1 capsid uncoating initiates after the first strand transfer of reverse transcription. Retrovirology, 2016. 13(1): p. 58.

61. Rankovic, S., et al., Reverse Transcription Mechanically Initiates HIV-1 Capsid Disassembly. J Virol, 2017. 91(12).

62. Yang, Y., T. Fricke, and F. Diaz-Griffero, Inhibition of reverse transcriptase activity increases stability of the HIV-1 core. J Virol, 2013. 87(1): p. 683–7.

63. Kessl, J.J., et al., HIV-1 Integrase Binds the Viral RNA Genome and Is Essential during Virion Morphogenesis. Cell, 2016. 166(5): p. 1257–1268 e12.

64. Fontana, J., et al., Distribution and Redistribution of HIV-1 Nucleocapsid Protein in Immature, Mature, and Integrase-Inhibited Virions: a Role for Integrase in Maturation. J Virol, 2015. 89(19): p. 9765–80.

65. Jurado, K.A., et al., Allosteric integrase inhibitor potency is determined through the inhibition of HIV-1 particle maturation. Proc Natl Acad Sci U S A, 2013. 110(21): p. 8690–5.

66. Engelman, A., In vivo analysis of retroviral integrase structure and function. Adv Virus Res, 1999. 52: p. 411–26.

67. Leavitt, A.D., et al., Human immunodeficiency virus type 1 integrase mutants retain in vitro integrase activity yet fail to integrate viral DNA efficiently during infection. J Virol, 1996. 70(2): p. 721–8.

68. Lu, R., H.Z. Ghory, and A. Engelman, Genetic analyses of conserved residues in the carboxyl-terminal domain of human immunodeficiency virus type 1 integrase. J Virol, 2005. 79(16): p. 10356–68.

69. Lu, R., et al., Class II integrase mutants with changes in putative nuclear localization signals are primarily blocked at a postnuclear entry step of human immunodeficiency virus type 1 replication. J Virol, 2004. 78(23): p. 12735–46.

70. Nakamura, T., et al., Lack of infectivity of HIV-1 integrase zinc finger-like domain mutant with morphologically normal maturation. Biochem Biophys Res Commun, 1997. 239(3): p. 715–22.

71. Shin, C.G., et al., Genetic analysis of the human immunodeficiency virus type 1 integrase protein. J Virol, 1994. 68(3): p. 1633–42.

72. Wu, X., et al., Human immunodeficiency virus type 1 integrase protein promotes reverse transcription through specific interactions with the nucleoprotein reverse transcription complex. J Virol, 1999. 73(3): p. 2126–35.

73. Ao, Z., et al., Contribution of the C-terminal tri-lysine regions of human immunodeficiency virus type 1 integrase for efficient reverse transcription and viral DNA nuclear import. Retrovirology, 2005. 2: p. 62.

74. Busschots, K., et al., Identification of the LEDGF/p75 binding site in HIV-1 integrase. J Mol Biol, 2007. 365(5): p. 1480–92.

75. Engelman, A., et al., Structure-based mutagenesis of the catalytic domain of human immunodeficiency virus type 1 integrase. J Virol, 1997. 71(5): p. 3507–14.

76. Limon, A., et al., Nuclear localization of human immunodeficiency virus type 1 preintegration complexes (PICs): V165A and R166A are pleiotropic integrase mutants primarily defective for integration, not PIC nuclear import. J Virol, 2002. 76(21): p. 10598–607.

77. Lloyd, A.G., et al., Characterization of HIV-1 integrase N-terminal mutant viruses. Virology, 2007. 360(1): p. 129–35.

78. Lu, R., et al., Lys-34, dispensable for integrase catalysis, is required for preintegration complex function and human immunodeficiency virus type 1 replication. J Virol, 2005. 79(19): p. 12584–91.

79. Masuda, T., et al., Genetic analysis of human immunodeficiency virus type 1 integrase and the U3 att site: unusual phenotype of mutants in the zinc finger-like domain. J Virol, 1995. 69(11): p. 6687–96.

80. Rahman, S., et al., Structure-based mutagenesis of the integrase-LEDGF/p75 interface uncouples a strict correlation between in vitro protein binding and HIV-1 fitness. Virology, 2007. 357(1): p. 79–90.

81. Riviere, L., J.L. Darlix, and A. Cimarelli, Analysis of the viral elements required in the nuclear import of HIV-1 DNA. J Virol, 2010. 84(2): p. 729–39.

82. Tsurutani, N., et al., Identification of critical amino acid residues in human immunodeficiency virus type 1 IN required for efficient proviral DNA formation at steps prior to integration in dividing and nondividing cells. J Virol, 2000. 74(10): p. 4795–806.

83. Wiskerchen, M. and M.A. Muesing, Human immunodeficiency virus type 1 integrase: effects of mutations on viral ability to integrate, direct viral gene expression from unintegrated viral DNA templates, and sustain viral propagation in primary cells. J Virol, 1995. 69(1): p. 376–86.

84. Zhu, K., C. Dobard, and S.A. Chow, Requirement for integrase during reverse transcription of human immunodeficiency virus type 1 and the effect of cysteine mutations of integrase on its interactions with reverse transcriptase. J Virol, 2004. 78(10): p. 5045–55.

85. De Houwer, S., et al., The HIV-1 integrase mutant R263A/K264A is 2-fold defective for TRN-SR2 binding and viral nuclear import. J Biol Chem, 2014. 289(36): p. 25351–61.

86. Johnson, B.C., et al., A homology model of HIV-1 integrase and analysis of mutations designed to test the model. J Mol Biol, 2013. 425(12): p. 2133–46.

87. Mohammed, K.D., M.B. Topper, and M.A. Muesing, Sequential deletion of the integrase (Gag-Pol) carboxyl terminus reveals distinct phenotypic classes of defective HIV-1. J Virol, 2011. 85(10): p. 4654–66.

88. Shehu-Xhilaga, M., et al., The conformation of the mature dimeric human immunodeficiency virus type 1 RNA genome requires packaging of pol protein. J Virol, 2002. 76(9): p. 4331–40.

89. Engelman, A., et al., Multiple effects of mutations in human immunodeficiency virus type 1 integrase on viral replication. J Virol, 1995. 69(5): p. 2729–36.

90. Madison, M.K., et al., Allosteric HIV-1 Integrase Inhibitors Lead to Premature Degradation of the Viral RNA Genome and Integrase in Target Cells. J Virol, 2017. 91(17).

91. Wang, W., et al., Inhibition of HIV-1 Maturation via Small-Molecule Targeting of the Amino-Terminal Domain in the Viral Capsid Protein. J Virol, 2017. 91(9).

92. Da Silva Santos, C., K. Tartour, and A. Cimarelli, A Novel Entry/Uncoating Assay Reveals the Presence of at Least Two Species of Viral Capsids During Synchronized HIV-1 Infection. PLoS Pathog, 2016. 12(9): p. e1005897.

93. Aiken, C., Cell-free assays for HIV-1 uncoating. Methods Mol Biol, 2009. 485: p. 41–53.

94. Yang, Y., J. Luban, and F. Diaz-Griffero, The fate of HIV-1 capsid: a biochemical assay for HIV-1 uncoating. Methods Mol Biol, 2014. 1087: p. 29–36.

95. Kutluay, S.B., D. Perez-Caballero, and P.D. Bieniasz, Fates of retroviral core components during unrestricted and TRIM5-restricted infection. PLoS Pathog, 2013. 9(3): p. e1003214.

96. Goudreau, N., et al., Novel inhibitor binding site discovery on HIV-1 capsid N-terminal domain by NMR and X-ray crystallography. ACS Chem Biol, 2013. 8(5): p. 1074–82.

97. Lemke, C.T., et al., A novel inhibitor-binding site on the HIV-1 capsid N-terminal domain leads to improved crystallization via compound-mediated dimerization. Acta Crystallogr D Biol Crystallogr, 2013. 69(Pt 6): p. 1115–23.

98. Kutluay, S.B. and P.D. Bieniasz, Analysis of HIV-1 Gag-RNA Interactions in Cells and Virions by CLIP-seq. Methods Mol Biol, 2016. 1354: p. 119–31.

99. Kutluay, S.B., et al., Global changes in the RNA binding specificity of HIV-1 gag regulate virion genesis. Cell, 2014. 159(5): p. 1096–109.

100. Douglas, C.C., et al., Investigation of N-terminal domain charged residues on the assembly and stability of HIV-1 CA. Biochemistry, 2004. 43(32): p. 10435–41.

101. Yang, R., et al., Second-site suppressors of HIV-1 capsid mutations: restoration of intracellular activities without correction of intrinsic capsid stability defects. Retrovirology, 2012. 9: p. 30.

102. Welker, R., et al., Biochemical and structural analysis of isolated mature cores of human immunodeficiency virus type 1. J Virol, 2000. 74(3): p. 1168–77.

103. Shi, J., et al., Compensatory substitutions in the HIV-1 capsid reduce the fitness cost associated with resistance to a capsid-targeting small-molecule inhibitor. J Virol, 2015. 89(1): p. 208–19.

104. Siddiqui, M.A., et al., A Novel Phenotype Links HIV-1 Capsid Stability to cGAS-Mediated DNA Sensing. J Virol, 2019. 93(16).

105. Fricke, T., et al., Human cytosolic extracts stabilize the HIV-1 core. J Virol, 2013. 87(19): p. 10587–97.

106. Ganser-Pornillos, B.K., A. Cheng, and M. Yeager, Structure of full-length HIV-1 CA: a model for the mature capsid lattice. Cell, 2007. 131(1): p. 70–9.

107. Shi, J. and C. Aiken, Saturation of TRIM5 alpha-mediated restriction of HIV-1 infection depends on the stability of the incoming viral capsid. Virology, 2006. 350(2): p. 493–500.

108. Mulder, L.C. and M.A. Muesing, Degradation of HIV-1 integrase by the N-end rule pathway. J Biol Chem, 2000. 275(38): p. 29749–53.

109. Ali, H., et al., Cellular TRIM33 restrains HIV-1 infection by targeting viral integrase for proteasomal degradation. Nat Commun, 2019. 10(1): p. 926.

110. Llano, M., et al., Lens epithelium-derived growth factor/p75 prevents proteasomal degradation of HIV-1 integrase. J Biol Chem, 2004. 279(53): p. 55570–7.

111. Zheng, Y., et al., Host protein Ku70 binds and protects HIV-1 integrase from proteasomal degradation and is required for HIV replication. J Biol Chem, 2011. 286(20): p. 17722–35.

112. Devroe, E., A. Engelman, and P.A. Silver, Intracellular transport of human immunodeficiency virus type 1 integrase. J Cell Sci, 2003. 116(Pt 21): p. 4401–8.

113. Francis, A.C., et al., Second generation imaging of nuclear/cytoplasmic HIV-1 complexes. AIDS Res Hum Retroviruses, 2014. 30(7): p. 717–26.

114. Maldarelli, F., M.A. Martin, and K. Strebel, Identification of posttranscriptionally active inhibitory sequences in human immunodeficiency virus type 1 RNA: novel level of gene regulation. J Virol, 1991. 65(11): p. 5732–43.

115. Schwartz, S., et al., Mutational inactivation of an inhibitory sequence in human immunodeficiency virus type 1 results in Rev-independent gag expression. J Virol, 1992. 66(12): p. 7176–82.

116. Schwartz, S., B.K. Felber, and G.N. Pavlakis, Distinct RNA sequences in the gag region of human immunodeficiency virus type 1 decrease RNA stability and inhibit expression in the absence of Rev protein. J Virol, 1992. 66(1): p. 150–9.

117. Wu, X. and G. Brewer, The regulation of mRNA stability in mammalian cells: 2.0. Gene, 2012. 500(1): p. 10–21.

118. Gorelick, R.J., et al., Characterization of the block in replication of nucleocapsid protein zinc finger mutants from moloney murine leukemia virus. J Virol, 1999. 73(10): p. 8185–95.

119. Miyazaki, Y., et al., An RNA structural switch regulates diploid genome packaging by Moloney murine leukemia virus. J Mol Biol, 2010. 396(1): p. 141–52.

120. Sakuragi, J., T. Shioda, and A.T. Panganiban, Duplication of the primary encapsidation and dimer linkage region of human immunodeficiency virus type 1 RNA results in the appearance of monomeric RNA in virions. J Virol, 2001. 75(6): p. 2557–65.

121. Tsang, J., et al., HIV-1 infection of macrophages is dependent on evasion of innate immune cellular activation. AIDS, 2009. 23(17): p. 2255–63.

122. Yan, N., et al., The cytosolic exonuclease TREX1 inhibits the innate immune response to human immunodeficiency virus type 1. Nat Immunol, 2010. 11(11): p. 1005–13.

123. Altfeld, M. and M. Gale, Jr., Innate immunity against HIV-1 infection. Nat Immunol, 2015. 16(6): p. 554–62.

124. Jonsson, K.L., et al., IFI16 is required for DNA sensing in human macrophages by promoting production and function of cGAMP. Nat Commun, 2017. 8: p. 14391.

125. Yoh, S.M., et al., PQBP1 Is a Proximal Sensor of the cGAS-Dependent Innate Response to HIV-1. Cell, 2015. 161(6): p. 1293–1305.

126. Blair, W.S., et al., HIV capsid is a tractable target for small molecule therapeutic intervention. PLoS Pathog, 2010. 6(12): p. e1001220.

127. Sticht, J., et al., A peptide inhibitor of HIV-1 assembly in vitro. Nat Struct Mol Biol, 2005. 12(8): p. 671–7.

128. Tang, C., et al., Antiviral inhibition of the HIV-1 capsid protein. J Mol Biol, 2003. 327(5): p. 1013–20.

129. Yant, S.R., et al., A highly potent long-acting small-molecule HIV-1 capsid inhibitor with efficacy in a humanized mouse model. Nat Med, 2019. 25(9): p. 1377–1384.

130. Shi, J., et al., Small-molecule inhibition of human immunodeficiency virus type 1 infection by virus capsid destabilization. J Virol, 2011. 85(1): p. 542–9.

131. Lamorte, L., et al., Discovery of novel small-molecule HIV-1 replication inhibitors that stabilize capsid complexes. Antimicrob Agents Chemother, 2013. 57(10): p. 4622–31.

132. Thenin-Houssier, S., et al., Ebselen, a Small-Molecule Capsid Inhibitor of HIV-1 Replication. Antimicrob Agents Chemother, 2016. 60(4): p. 2195–208.

133. Kortagere, S., et al., Inhibiting early-stage events in HIV-1 replication by small-molecule targeting of the HIV-1 capsid. J Virol, 2012. 86(16): p. 8472–81.

134. Bhattacharya, A., et al., Structural basis of HIV-1 capsid recognition by PF74 and CPSF6. Proc Natl Acad Sci U S A, 2014. 111(52): p. 18625–30.

135. Price, A.J., et al., Host cofactors and pharmacologic ligands share an essential interface in HIV-1 capsid that is lost upon disassembly. PLoS Pathog, 2014. 10(10): p. e1004459.

136. Fricke, T., et al., BI-2 destabilizes HIV-1 cores during infection and Prevents Binding of CPSF6 to the HIV-1 Capsid. Retrovirology, 2014. 11: p. 120.

137. Rihn, S.J., et al., Extreme genetic fragility of the HIV-1 capsid. PLoS Pathog, 2013. 9(6): p. e1003461.

138. Zennou, V., et al., APOBEC3G incorporation into human immunodeficiency virus type 1 particles. J Virol, 2004. 78(21): p. 12058–61.

139. Cowan, S., et al., Cellular inhibitors with Fv1-like activity restrict human and simian immunodeficiency virus tropism. Proc Natl Acad Sci U S A, 2002. 99(18): p. 11914–9.

140. Hatziioannou, T., et al., Restriction of multiple divergent retroviruses by Lv1 and Ref1. EMBO J, 2003. 22(3): p. 385–94.

141. Bouyac-Bertoia, M., et al., HIV-1 infection requires a functional integrase NLS. Mol Cell, 2001. 7(5): p. 1025–35.

142. Pizzato, M., et al., A one-step SYBR Green I-based product-enhanced reverse transcriptase assay for the quantitation of retroviruses in cell culture supernatants. J Virol Methods, 2009. 156(1-2): p. 1–7.

143. Yoo, S., et al., Molecular recognition in the HIV-1 capsid/cyclophilin A complex. J Mol Biol, 1997. 269(5): p. 780–95.

144. Gres, A.T., et al., STRUCTURAL VIROLOGY. X-ray crystal structures of native HIV-1 capsid protein reveal conformational variability. Science, 2015. 349(6243): p. 99–103.

